# Left-Right Asymmetry in the Breast TME is Associated with Distinct Bioelectric and Epigenetic Tumor Features

**DOI:** 10.1101/2025.06.30.662430

**Authors:** Sebastián Real, Sergio Laurito, Nadia Bannoud, Pablo Gonzalez, Oscar Bello, Diego Croci, Joao Carvalho, María Roqué

## Abstract

Breast cancer heterogeneity is shaped not only by intrinsic tumor properties but by the surrounding microenvironment (TME). Here we report a left–right (L–R) asymmetry in the breast TME associated to tumor features. Bioinformatic analysis of TCGA RNA-seq and histological data revealed that R-sided tumors display higher stromal scores and are cancer-associated fibroblasts (CAFs) signatures. This difference was validated in paired L–R PDX and xenograft models, in which R-sided tumors showed increased α-SMA expression. The asymmetry extended to tumor bioelectricity: across PDX and xenograft models, and in cells cultured with L-or R-conditioned media, L-sided tumors consistently displayed a more depolarized membrane potential than their R counterparts. In vitro, this bioelectric difference was methylation-dependent — abolished by the DNA methyltransferase inhibitor decitabine — and reversible when the laterality of the conditioned media was inverted. Methylome analysis of animal models identified the gap-junction gene *GJA1* as significantly hypermethylated in L-sided tumors, accompanied by reduced expression of connexin genes. A computational model showed that membrane potential and methylation form two stable states, depending on gap-junction coupling strength. Together, these findings identify L–R asymmetry as an overlooked layer of breast tumor heterogeneity that integrates stromal, bioelectric, and epigenetic features. This framework may extend to other paired organs and suggests that tumor laterality could have biological and therapeutic relevance.

## INTRODUCTION

Breast cancer is the most frequently diagnosed malignancy and a leading cause of cancer-related mortality in women worldwide^1^. Despite major advances in early detection and targeted therapies, it remains a substantial global health burden^1^. A defining characteristic of breast cancer is its marked heterogeneity, which underlies differences in disease progression, prognosis, and therapeutic response^2^. Historically, this heterogeneity has been studied primarily through intrinsic tumor features, including molecular subtypes, genomic alterations, and transcriptional profiles^3^. However, increasing evidence indicates that extrinsic factors within the tumor microenvironment (TME) are equally important determinants of tumor behavior^4^. Stromal composition, immune-cell infiltration, extracellular matrix remodeling, and metabolic interactions can profoundly influence cancer progression and treatment response^4^. While significant progress has been made in classifying intrinsic tumor subtypes, the contribution of anatomical and physiological context remains comparatively underexplored.

Among these extrinsic influences, anatomical laterality represents a largely overlooked source of variation. A growing body of evidence, including findings from our group and others^5–11^, suggests that left-right (L-R) asymmetry may contribute to breast cancer heterogeneity. Epidemiological studies consistently report a modest but reproducible predominance of L-sided breast cancer, with an incidence ratio of approximately 1.15 compared with R-sided ones^5,12^. Beyond incidence, breast tumors of different sides exhibit distinct biological characteristics^13^, and tumor laterality has gained clinical relevance following reports that L-sided breast cancers display more aggressive features and poorer outcomes than the R-sided counterparts^7^. Similar asymmetries in cancer incidence have been documented in other bilateral organs, including the lung, testis, and ovary^5^, suggesting that paired organs may not provide identical contexts for tumor development.

This possibility is supported by fundamental principles of developmental biology. L-R asymmetry is established early during embryogenesis through conserved signaling pathways, including Nodal, Lefty, and Pitx2^14,15^, which generate persistent differences in organ structure and physiology. Beyond obvious anatomical asymmetries, subtle side-specific variations have been described in vascular drainage, lymphatic circulation, neural regulation, and immune function^16–18^. These observations indicate that the L and R sides of the body constitute distinct biological environments. Such asymmetries have been implicated in disease susceptibility, including neurological disorders characterized by unilateral manifestations^19^. In cancer, laterality has also been associated with biological and clinical differences. For example,R-sided colorectal cancers exhibit distinct molecular profiles and poorer prognosis than L-sided tumors^20^, while asymmetries in lung structure and physiology have been proposed to influence tumor progression^21^. Nevertheless, for cancers arising in bilateral organs, the mechanisms linking normal physiological asymmetry to tumor heterogeneity remain poorly understood.

Breast cancer provides a particularly attractive system in which to investigate this question. Our previous studies demonstrated that L- and R-sided breast tumors differ at multiple biological levels, including DNA methylation patterns, membrane potential (Vmem), ion-channel (ICH) gene expression, and proliferative activity^9–11^. These findings suggest that laterality may influence fundamental regulatory processes within tumors. Studies have demonstrated that L-R mammary glands differ in baseline molecular composition, are independently regulated during normal and neoplastic development, and generate distinct gene-expression signatures^18^. These observations raise the possibility that breast tumors arise within biologically distinct L and R microenvironments that may differentially shape tumor evolution.

Bioelectric signaling^22^ and epigenetic regulation^23^ are increasingly recognized as important contributors to tumor heterogeneity. Both participate in cell-to-cell communication and in the dynamic interactions between tumor cells and their microenvironment. In parallel, the TME is now understood as an active regulator of cancer progression rather than a passive scaffold. Stromal components such as cancer-associated fibroblasts (CAFs) remodel the extracellular matrix, secrete growth factors and cytokines, promote angiogenesis, and influence immune responses and tumor-cell behavior^24^.

We therefore hypothesized that anatomical laterality establishes distinct TMEs that may be associated with tumor bioelectric states through epigenetic regulation of ion-channel genes. To test this hypothesis, we combined bioinformatic analyses of human breast cancers with histological validation, in vivo tumor models, methylome profiling, and computational modeling to determine whether stromal, bioelectric, and epigenetic asymmetries converge as interconnected features of L-R-sided breast tumors.

## RESULTS

### 1. R-SIDED BREAST TUMORS EXHIBIT STROMAL ENRICHMENT

#### Stromal Scores are Enriched in R-Sided IDCs

To investigate potential bioinformatic differences in the tumor microenvironment (TME) of L and R breast tumors, we analyzed primary invasive ductal carcinomas (IDCs) from the TCGA-BRCA cohort obtained from The Cancer Genome Atlas (TCGA-BRCA). A total of 784 IDC samples were available for analysis.

Because stromal cells are major components of the TME, we initially aimed to focus on tumors with lower purity scores, reasoning that reduced tumor purity may reflect greater stromal representation and thereby increase sensitivity to detect TME-related differences. We proposed for this the Consensus Purity Estimation score, CPE, which estimates tumor purity based on RNA-Seq data from tumoral and non-cancerous cells^28^. To determine whether CPE reliably captured stromal abundance, we validated the metric using two independent approaches. First, we analyzed 20 H&E-stained whole-slide images from TCGA breast IDCs, selecting the 10 tumors with the highest and the 10 with the lowest RNA-seq-derived stromal scores to maximize contrast in microenvironment composition (Clinical Data in Supplementary Data 1). Relative stromal abundance [connective cells / (neoplastic + connective cells)] was quantified using <u>HoVer-Net</u> ((https://github.com/vqdang/hover_net), a multi-branch network for nuclear instance segmentation and classification^25^) and correlated with CPE scores. A significant inverse correlation was found (Pearson r = −0.519, p = 0.018), indicating that lower CPE values correspond indeed to higher stromal content (Supplementary Figure 1a and Supplementary Data 2). Second, stromal abundance was estimated across the complete IDC cohort using <u>xCell</u>. This analysis also confirmed a significant inverse association between stromal content and CPE (Pearson r = −0.39, p < 0.0001; Supplementary Figure 1b), supporting the use of CPE as an inverse surrogate of tumor stromal abundance.

Based on these validations, we further applied the CPE score and selected tumors with a CPE ≤70%, yielding a subset of 276 IDC cases which were separated to use as discovery set and in parallel the entire cohort was used as extension/validation cohort. The stromal and immune scores were calculated by xCell from transformed single-sample GSEA (ssGSEA) normalized RNA-Seq counts of the 276-sample. L vs R comparisons of the scores were performed using Rstats and the UCSC Xena Functional Genomics Explorer (https://xenabrowser.net/). The results revealed that R-sided tumors exhibited significantly higher stromal scores compared to L-sided tumors (Welch’s corrected t-test, p = 0.006, t = -2.734; Figure 1a). This difference persisted in the full 784-sample IDC cohort (Welch’s corrected t-test, p = 0.01, t = -2.398; Figure 1b). When stratifying by PAM50 subtypes, the differences did not reach statistical significance (Wilcoxon rank-sum test, Supplementary Figure 2a-b) even though the differences maintained similar directions (except for Basal or HER2 tumors, depending on the cohorts). Regarding the immune scores, no significant L-R differences were observed in either cohorts (Supplementary Figure 2c-d).

**Figure 1.**
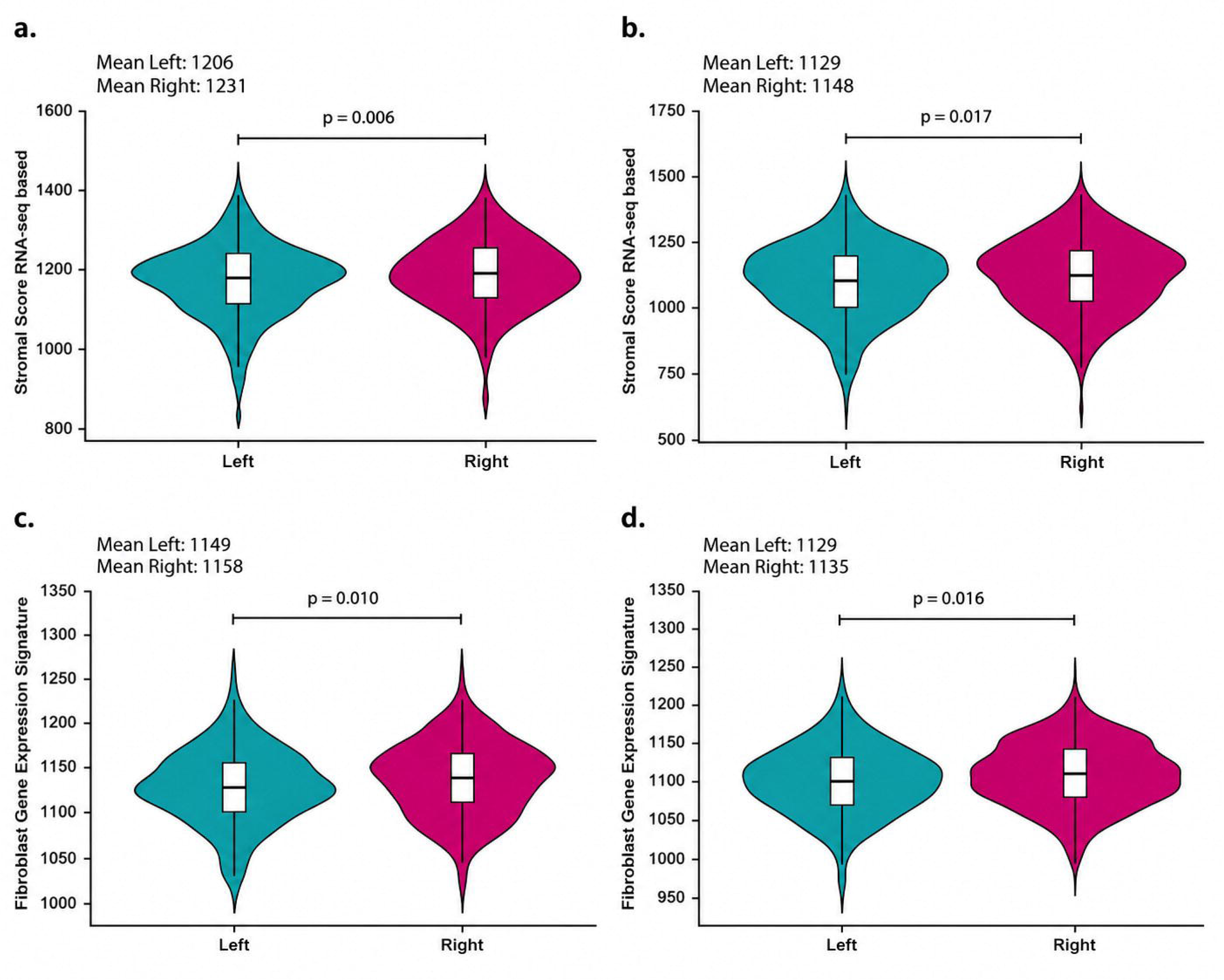
Stromal Composition in L-R IDCs. **a.** Stromal gene signature scores in 276 TCGA IDC samples (≤70% purity) (L-tumors= 153 and R-tumors=123) show significantly higher scores in R-tumors (Welch’s corrected t-test, p = 0.006). b. The same comparison in the full 784 IDC cohort (no purity restriction) (L-tumors=424 and R-tumors= 360) confirms higher stromal scores in R-tumors (Welch’s corrected t-test, p = 0.01). c. Fibroblast gene signature scores in 276 TCGA IDC samples (≤70% purity) are elevated in R-tumors (Welch’s corrected t-test, p = 0.01 for Fibroblast_FANTOM_1). D. Fibroblast gene signature scores in the full 784 IDC cohort (no purity restriction) remain significantly elevated in R-tumors (Welch’s corrected t-test, p = 0.01). Stroma Score Gene Signature and Fibroblast Gene Expression Signature data were obtained from Xena Browser (https://xenabrowser.net/). The data are log₂-transformed and plotted on a linear axis. Plots were generated using ggplot2 package. Visualization was performed with Adobe Illustrator 2024.

#### Fibroblasts are Enriched in R-Sided IDCs

To further characterize the stromal differences, we analyzed bioinformatically the gene expression signatures of stromal cell subtypes, including fibroblasts, endothelial cells, adipocytes, and others. In the 276-sample cohort, fibroblasts were significantly more abundant in R-sided IDCs (Welch’s corrected t-test, p < 0.05), for the signatures extracted from xCell algorithm, including Fibroblast_FANTOM_1 (126 genes, p = 0.01, t = -2.528), Fibroblast_HPCA_1 (34 genes, p = 0.05), and Fibroblast_HPCA_2 (61 genes, p = 0.04) (Figure 1c; gene lists in Supplementary Data 3). These patterns were consistent in the full 784-sample IDC cohort (p = 0.01, Figure 1d). The R-sided increment of CAFs was observed across most of the PAM50 breast tumor subtypes although not reaching statistical significance (Supplementary Figure 2e-f).

To further investigate which CAF subtype was involved, we used single-cell RNA-Seq-derived gene signatures from Cords et al. (2023)^31^ for seven CAF subtypes (inflammatory [iCAF], dividing [dCAF], vascular, matrix, hsp, rare, and tumor CAFs (gene lists in Supplementary Data 4). No differences were found among the 276-sample cohort (p > 0.05). However, when expanding to the complete cohort, R-sided tumors exhibited higher iCAF expression (Unpaired t-test, p = 0.003), while L-sided tumors showed elevated dCAF expression (Unpaired t-test, p = 0.0026; Figure 2). The L-R differences of dCAFs and iCAFs were directionally similar across most PAM50 breast tumor subtypes but did not reach statistical significance (Supplementary Figure 2g).

**Figure 2.**
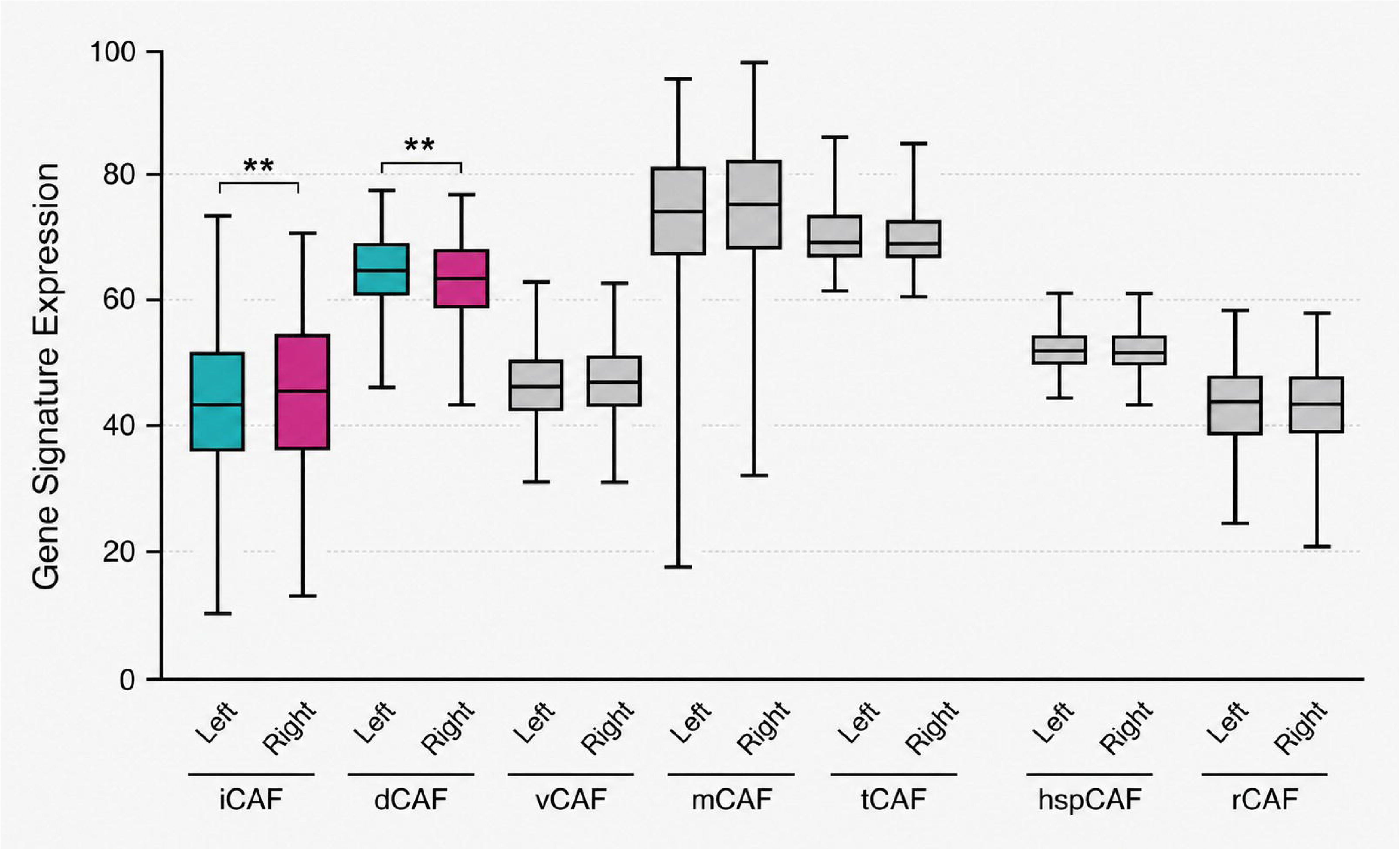
CAF Subtype Composition in L-R IDCs. L-R comparison of CAF subtype gene signatures in the 784 TCGA IDC cohort shows increased iCAF expression in R-tumors (Unpaired T-test, p = 0.003) and dCAF expression in L-tumors (Unpaired T-test, p = 0.0026). Blue and pink boxes indicate L-R significant differences, whereas gray boxes indicate no significant L-R differences. Data are presented as mean ± SEM. Analysis performed with GraphPad Prism v5, visualized with Adobe Illustrator 2024.

#### Histological Validation of Stromal Enrichment in R-sided Tumors

To validate the L-R stromal differences at tissue level, we analyzed bioinformatically the stromal component on 20 H&E-stained whole-slide images of breast IDCs from the TCGA dataset blinded to tumor laterality and outcome. To maximize contrast in stromal composition, all available IDC samples were ranked according to their transcriptome-derived stromal abundance estimates, and the 10 tumors with the highest and the 10 with the lowest stromal scores were selected. For each slide, regions of interest were defined, prioritizing nuclear density and technical staining quality (Methods). To minimize confounding factors, we confirmed no significant differences in age, tumor subtype, stage, size, or menopausal status between L and R groups (Supplementary Data 5). The relative stromal component was quantified as mentioned above (Connective cells/[Neoplastic cells + Connective cells]). R-sided tumors displayed a significantly higher stromal relative abundance (median: 0.395, IQR 0.238-0.601) compared to L-sided tumors (median: 0.108, IQR 0.0372-0.157], (Wilcoxon test, p = 0.002; Figure 3). By this, the histological evidence supported the RNA-Seq based bioinformatic findings.

**Figure 3.**
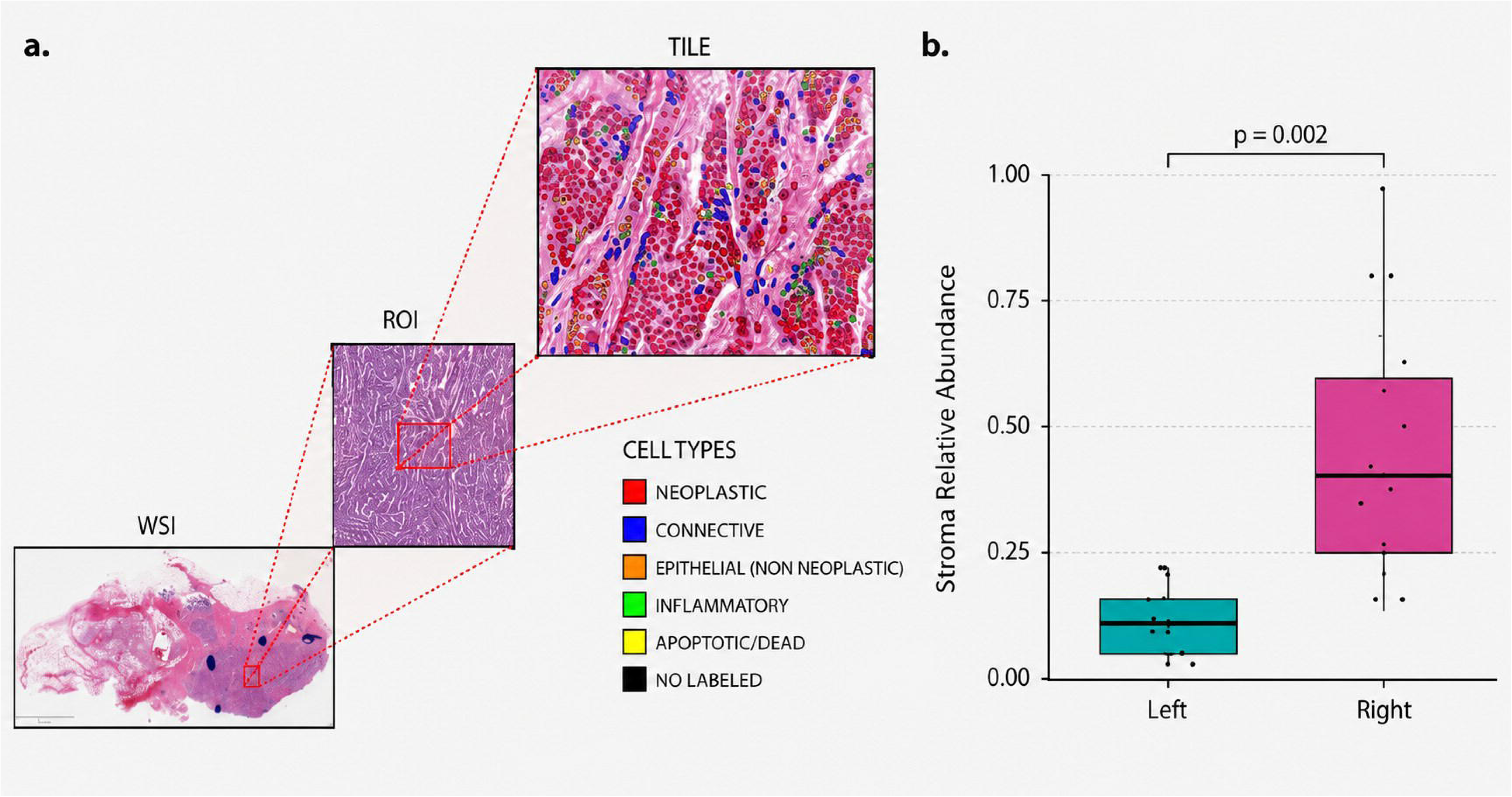
Stromal Fraction in L-R IDCs from TCGA H&E Images. **a.** Representative image of simultaneous nuclei segmentation and classification in H&E-stained tissue slides from the TCGA-BRCA project using HoVer-Net^29^. For each whole-slide image (WSI), a square region of interest (ROI, 2000 µm × 2000 µm) selected based on predefined technical quality criteria (e.g., staining consistency, absence of artifacts) while blinded to tumor laterality and outcomes. Each ROI was subsequently divided into non-overlapping tiles of 512 px (≈ 260 TILES per ROI). All tiles were processed with HoVer-Net to perform nuclei segmentation and classification. The output consisted of (i) an image with nuclei labeled by cell type (see image for color code) and (ii) a comma-separated values (CSV) file reporting the counts of each cell type per tile. **b**. Quantification of stromal component counts (relative abundance: connective tissue/neoplastic+connective ratio) in 10 L and 10 R TCGA IDC H&E slides shows higher stromal content in R-tumors (Wilcoxon test, p = 0.002). Data are presented as mean ± SEM. Analysis performed with digital pathology software (Hovernet), visualized with Adobe Illustrator 2024.

#### Increased α-SMA expression in R-Sided PDX and Xenograft Tumors

We next examined stromal myofibroblast-like cells in two complementary human breast cancer xenograft models: a patient-derived xenograft (PDX) and an MDA-MB-231 triple-negative breast cancer xenograft. As expected for human xenograft systems, the tumor stromal compartment, including fibroblasts, was of murine origin. αSMA is widely used in breast tumor stroma as a marker of activated myofibroblast-like stromal cells, which constitute the predominant αSMA-positive stromal population in breast tumors^26–28^, although is not exclusively expressed by cancer-associated fibroblasts. In these xenograft models the other α-SMA-positive populations are either absent or spatially restricted — myoepithelial cells are not reconstituted in MDA-MB-231 or PDX tumors, and pericytes and vascular smooth-muscle cells are confined to the perivascular compartment — so they are unlikely to account for the L–R differences observed. Three mice bearing bilateral PDX tumors (∼1000 mm³) were sacrificed, and αSMA-positive stromal signal was quantified by immunofluorescence and confocal microscopy using an Alexa-conjugated anti-α-smooth muscle actin (αSMA) antibody. To account for inter-animal variability, all αSMA/Hoechst values were normalized to the mean value of the corresponding R-sided tumor within each mouse. Consistent with the bioinformatic predictions, in all analyzed animals R-sided PDX tumors exhibited higher αSMA signal than their paired L-sided counterparts, reaching statistical significance in two of three mice (Mouse 1, p = 0.001; Mouse 2, p = 0.005; Mouse 3, p = 0.108; Figure 4a). Remarkably, the same pattern was reproduced in the MDA-MB-231 xenograft model, where R-sided tumors consistently displayed higher αSMA fluorescence than L-sided tumors across all four animals examined reaching significance in three of four (Mouse 1, p = 0.0003; Mouse 2, p = 0.0002; Mouse 3, p = 0.0021; Mouse 4, p = 0.16; Figure 4b). Representative immunofluorescence images from three L-sided and three R-sided tumors are shown in Figure 4c. Although the number of biological replicates was limited, the concordant direction observed across both independent models provides in vivo support for an enrichment of α-SMA-positive stromal cells in R-sided breast tumors.

**Figure 4.**
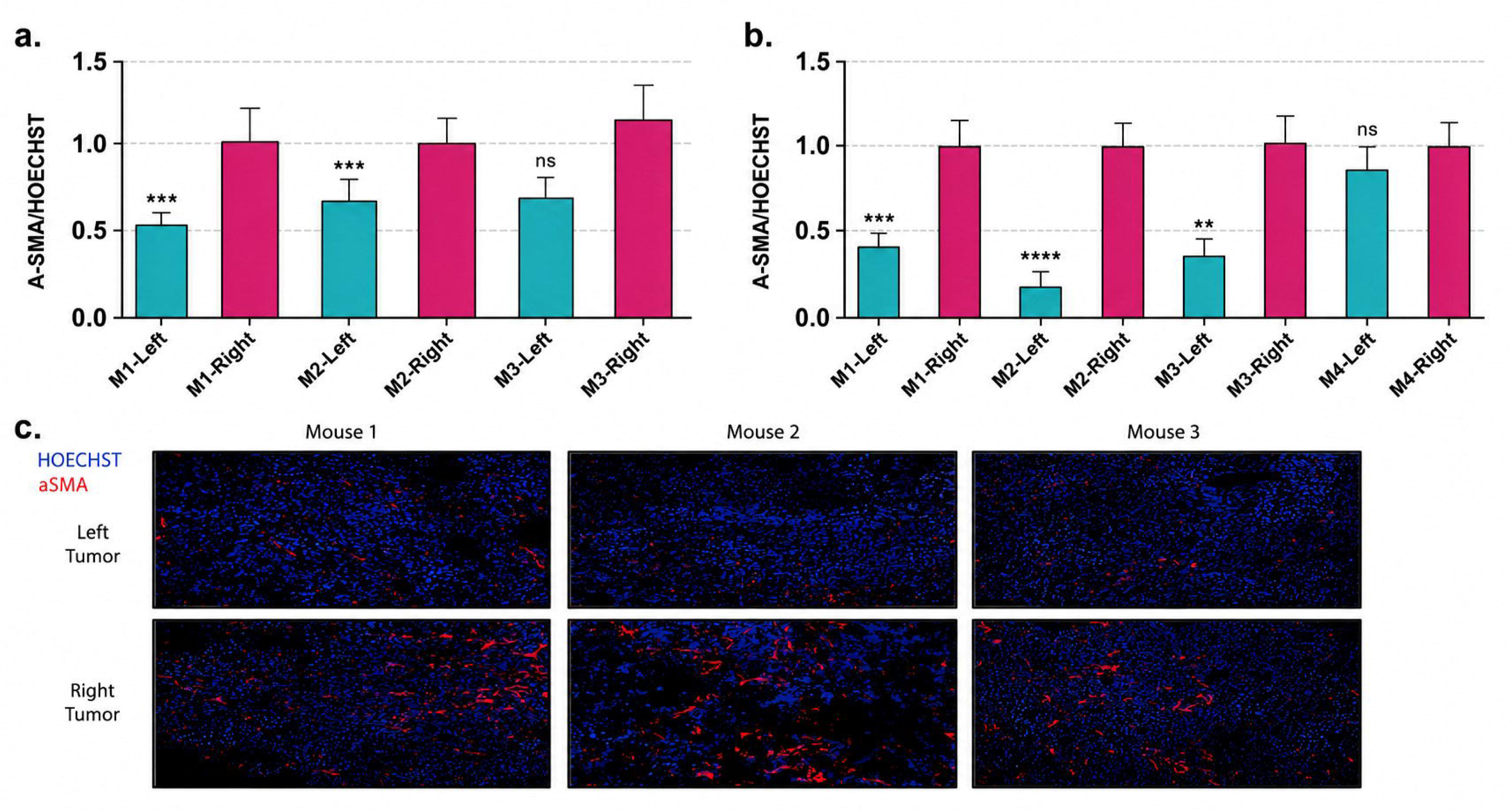
Increased α-SMA in R-Sided PDX and Xenograft Tumors. At least five ROIs were analyzed per tumor section from each bilateral PDX and MDA-MB-231 xenograft tumor by confocal microscopy. αSMA/Hoechst fluorescence values were normalized to the mean value of the corresponding R-sided tumor within each mouse to account for inter-animal variability. ROIs were used as technical replicates to estimate tumor-level signal intensity, whereas biological comparisons were performed at the mouse level. Across both models, R-sided tumors consistently exhibited higher αSMA signal than paired L-sided tumors. **a,** Quantification of αSMA/Hoechst fluorescence ratios in three bilateral PDX tumors (one-sample t-test; Mouse 1, *p* = 0.001; Mouse 2, *p* = 0.005; Mouse 3, *p* = 0.108 after outlier removal). **b,** Quantification of αSMA/Hoechst fluorescence ratios in four bilateral MDA-MB-231 xenografts (one-sample t-test; Mouse 1, *p* = 0.0003; Mouse 2, *p* = 0.0002; Mouse 3, *p* = 0.0021; Mouse 4, *p* = 0.16). Data are shown as mean ± SEM. **c,** Representative confocal images of MDA-MB-231 xenograft tumors (three L-sided and three R-sided tumors from three independent mice). Nuclei were stained with Hoechst (blue) and αSMA is shown in red.

#### Stromal Component Comparison in L-R PDX and Xenograft Tumors

To further extend the analysis to endothelial and immune cell populations, we performed flow cytometry assays on single-cell suspensions obtained from three PDX and four MDA-MB-231 xenograft pairs (L-R tumors, ∼1000 mm³). We quantified cell populations with specific markers for different immune cell types, endothelial cells and fibroblasts. Due to limited sample availability, not all animals were analyzed for every marker; however, at least three paired tumors (PDX and/or MDA-MB-231 xenografts) were included in each analysis. We found no significant differences in any of the immune cell populations (total leukocytes – CD45^+^ [Figure 5a], myeloid cells – CD11b^+^[Figure 5c], dendritic cells – CD11c^+^[Figure 5d]), or in endothelial cells – CD31^+^ (Figure 5b, right panel), either within individual models or when analyzed collectively (p > 0.05, Paired t-test). These findings are consistent with our bioinformatic xCell predictions. The CAF marker PDGFRα showed higher mean expression in R-sided tumors, although the difference was not statistically significant (mice n=5, paired *t*-test, p = 0.3; Figure 5b, left panel, framed in red).

**Figure 5.**
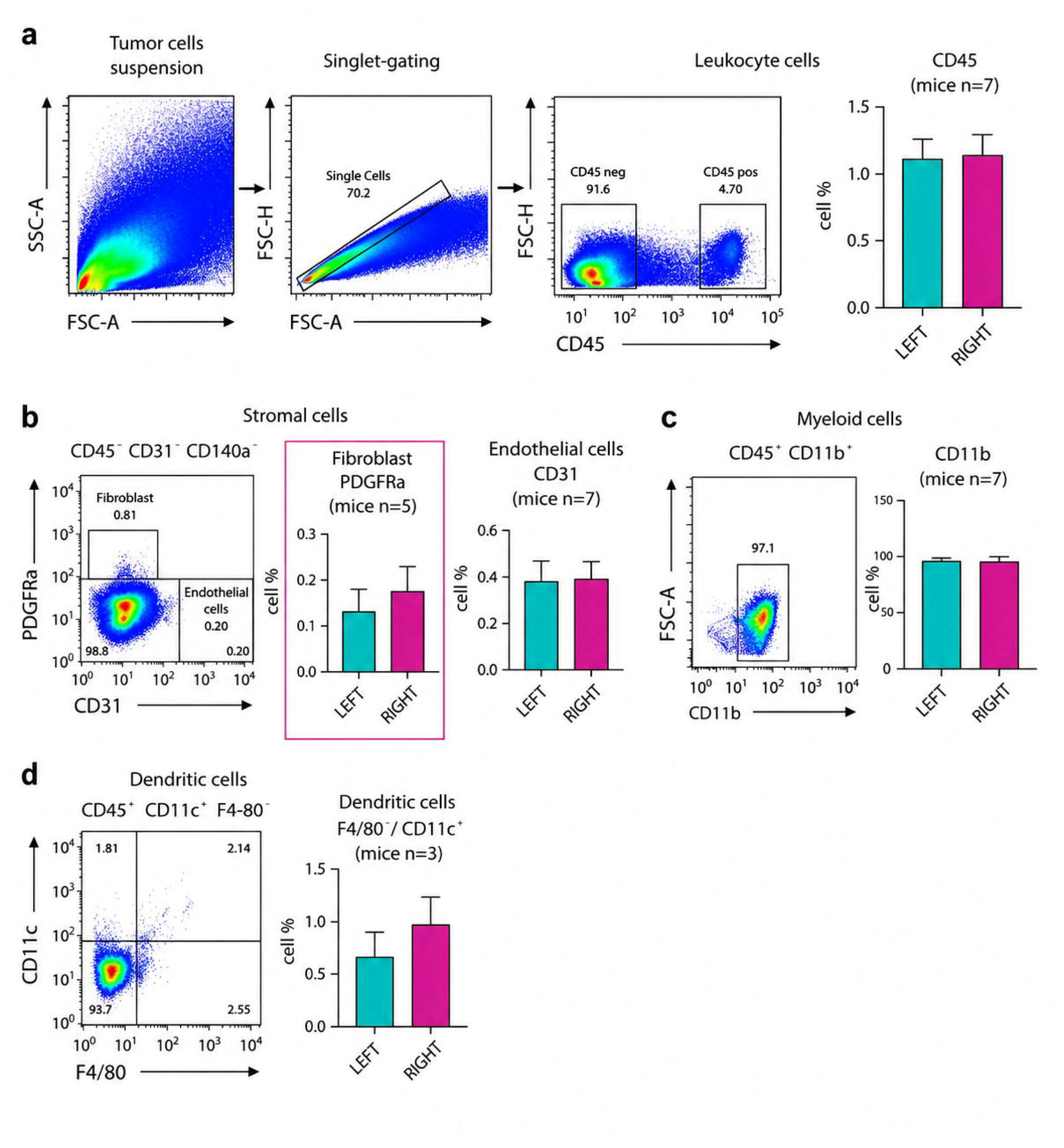
Flow Cytometry Analysis of L-R mice Tumors. **a**. Hierarchical gating strategy for tumor single-cell suspension to identify immune (CD45+) and non-immune stromal (CD45-) populations. FSC-A vs SSC-A: debris exclusion. FSC-H vs FSC-A: singlet gating. FSC-A vs CD45: discrimination of CD45⁺ immune cells and CD45⁻ non-immune stromal cells. From the CD45⁻ stromal fraction, (**b**) fibroblasts and endothelial cells were identified based on PDGFRα vs CD31 expression (fibroblasts: PDGFRα⁺CD31⁻; endothelial cells: CD31⁺). From the CD45⁺ immune fraction, (**c**) CD11b⁺ myeloid cells were gated, followed by (**d**) identification of dendritic cell populations based on CD11c vs F4/80. Representative plots are shown; quantitative data for each population are provided in the accompanying bar graphs. Fluorescence parameters were displayed using biexponential scaling, whereas forward and side scatter were plotted on a linear scale. Quantification of immune cell populations and endothelial cells shows no significant differences between L-R tumors (paired T-test, p > 0.05, Figure 5a, b-panel right, and c). Of the non-immune stromal cells, we highlight in red frame the higher mean PDGFRα expression in R-tumors, which did not reach statistical significance (mice: n=5, paired t-test, p = 0.3, Figure 5b-left panel). The apparent difference in dendritic cells (Figure 5d) was not considered reliable due to the limited number of analyzed animals (n=3 mice) and the large standard error. Data is presented as mean ± SEM. Analysis performed with FlowJo v10, visualized with Adobe Illustrator 2024.

So far, having established observations that suggest a lateralized stromal composition in both human cohorts and animal models, we next examined whether bioelectric properties of the tumors also differed by side.

### 2. L-R TME ASYMMETRY ASSOCIATES WITH DISTINCT TUMOR BIOELECTRICITY

#### L-sided tumors are depolarized in Xenografts and PDX models

We next aimed to test the hypothesis in clinically relevant models whether different stroma associate with tumor bioelectric differences. To this end, we used PDX and xenografted tumors. Unlike our previously studies on a spontaneous MMTV-PyMT model^11^, xenografting allows the implantation of genetically identical cells into both sides of the same host. This design controls initial genetic and epigenetic variability, ensuring that any bioelectric divergence arises from the local mammary gland environment rather than intrinsic properties of the inoculated cells. Thus, xenografts provide a rigorous framework to determine whether host tissue laterality can condition tumor bioelectric states.

We generated three xenograft mice by injecting equal numbers of triple negative human breast cancer cells MDA-MB-231 into the L and R fourth mammary glands. When measuring the Vmem of tumors using the ratio between two oppositely charged probes (Mitotracker -MT- / DiBAC -DB-), in all three cases we observed a consistent L– R asymmetry. L-sided tumors displayed significantly lower MT/DB fluorescence ratios when relativized to R, indicating a more depolarized state (mice n=3, Figure 6a; one-sample t-test against 1, t = 5.792, df = 2, p = 0.029). When normal mammary tissue was expressed relative to R-sided tumors, it did not differ significantly from 1 (one-sample t-test against 1, p = 0.376), indicating that R tumors maintain a Vmem close to that of homeostatic tissue. In contrast, the maximally depolarized KCl control differed markedly from R (one-sample t-test against 1, p < 0.0001). Together, these results indicate that R tumors adopt an electrical state close to normal tissue, while L tumors shift toward depolarization.

**Figure 6.**
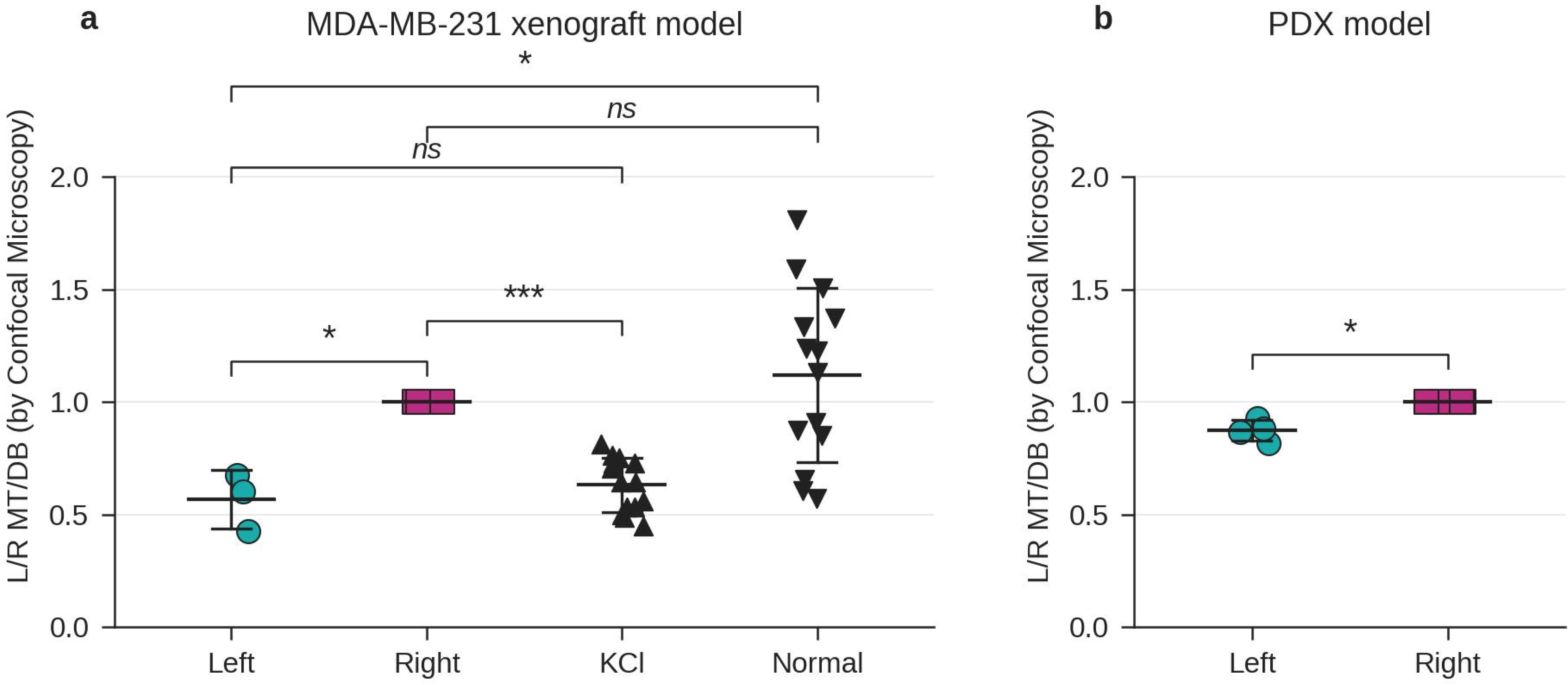
Vmem in L-R xenograft and PDX tumors. **a.** Tumor Vmem was assessed in thin tissue sections from 3 pairs of L- and R-sided xenograft tumors of MDA-MB231 cells using the MT/DB fluorescence ratio, quantified by confocal microscopy through summed Z-stack intensity projections. L-sided tumors exhibited significantly lower MT/DB ratios than their R-sided counterparts (one-sample t-test against 1, t = 5.792, df = 2, p = 0.029), indicating a more depolarized state. Normal mammary tissue, expressed relative to R, did not differ from 1 (one-sample t-test against 1, p = 0.376), whereas the maximally depolarized KCl control did (one-sample t-test against 1, p < 0.0001). All comparisons were performed as one-sample t-tests of the indicated ratio against a theoretical value of 1. **b**. Patient-derived xenografts (PDX) established from a single HER2-positive breast tumor were fragmented and implanted bilaterally into the L and R mammary glands of NSG mice, such that laterality denotes implantation site rather than donor-tumor side. In four paired tumors, L-sided PDX tumors displayed a significantly lower MT/DB ratio than their paired R-sided tumors (n = 4 mice; one-sample t-test of the L/R ratio against a theoretical value of 1: t = 5.49, df = 3, p = 0.012), confirming a consistent depolarization bias in L tumors across distinct breast cancer models. All comparisons in this figure were performed as two-tailed one-sample t-tests of the indicated ratio (L/R, NT/R, or KCl/R) against a theoretical value of 1.

We further extended our analyses to a patient-derived model. Importantly, as for the xenografts, laterality here refers to the implantation site: fragments of a single HER2+ PDX tumor were transplanted bilaterally into the L and R mammary glands of the same host, so any L-R divergence reflects the local gland environment rather than the donor tumor’s original side. In this PDX system, we examined four pairs of bilateral tumors and observed the same asymmetry: L-sided tumors displayed a significantly lower MT/DB ratio than their paired R-sided tumors (n = 4 mice; one-sample t-test of the L/R ratio against a theoretical value of 1: t = 5.49, df = 3, p = 0.012, Figure 6b). Thus, two independent *in vivo* models-xenografts and PDX-L tumors reproducibly adopt a depolarized state, whereas R tumors maintain an electrical profile close to that of normal tissue.

Together, these results show that the L-R bioelectric asymmetry is reproducible across genetically distinct tumor models and host environments. Considering the TME as a key modulator of this asymmetry, we next turned to cell culture systems to test the reversibility of the phenomenon.

#### Bioelectric States are Reversible and Epigenetically Regulated in cell culture

We have in previous publications demonstrated that extracts from human healthy breast tissues were capable of inducing L–R bioelectric shifts in cultured cells, consistently showing that extracts from the L breast depolarized cancer cells compared with extracts from the R side^6, 7^. In this work, we expanded the approach by testing extracts from healthy bovine mammary (udder) tissue applied to MDA-MB-231 cells, which produced the same L–R shift observed with healthy human breast-tissue extracts (Supplementary Figure 3). Although these experiments do not provide new mechanistic insights, they offer technical validation, establish bovine tissue as a scalable model system, and further support the concept that L–R microenvironmental differences can consistently modulate cellular Vmem.

To further assess the impact of the “side environment,” we examined whether the bioelectrical plasticity of cancer cells was reversible. In the conditioned-culture model, MDA-MB-231 cells were exposed for 5 days to healthy human breast-tissue extract from one side (L or R) and then switched to the opposite-side extract for a further 5 days (L→R and R→L; diagram in Figure 7a). As expected, cells initially exposed to L extract exhibited a lower MT/DB ratio relative to R-treated cells (Figure 7b, *Original extracts,* MT/DB <1). Upon switching from L to R extract, the MT/DB ratio increased (in some cases >1), indicating a shift toward a more polarized state (Figure 7b, *Inverted extracts*; unpaired t-test, p = 0.05). This suggests that the effect is reversible upon altering the extracellular environment.

**Figure 7.**
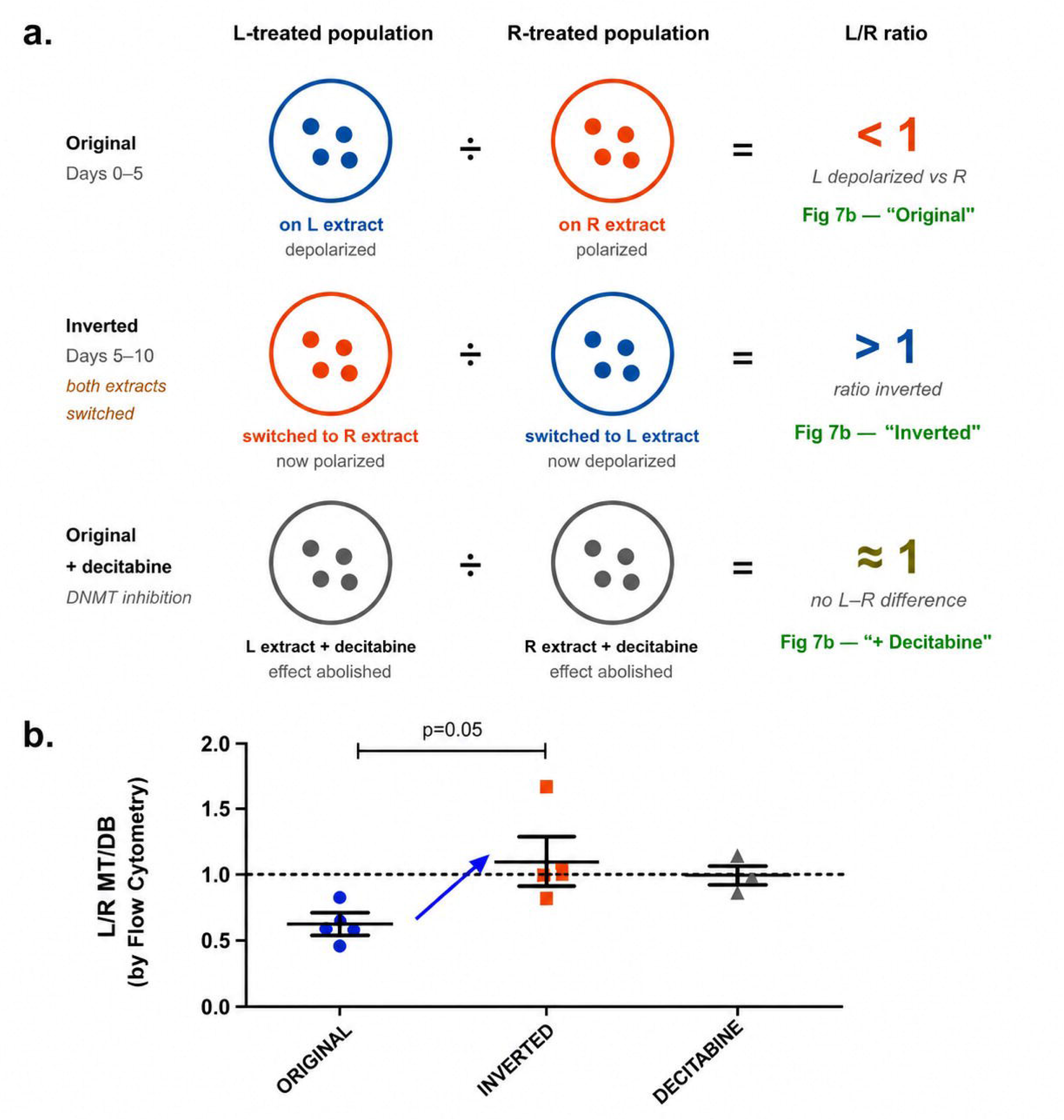
Reversibility and epigenetic dependence of L–R bioelectric shifts. **a.** Diagram of the conditioned-culture design. Two parallel populations of MDA-MB-231 cells were maintained on healthy human L- or R-breast extract, and the bioelectric asymmetry was expressed as the ratio of their Vmem (MT/DB of L-treated / MT/DB of R-treated cells). *Original*: L-treated cells are depolarized relative to R-treated cells (ratio < 1). *Inverted*: after switching both populations to the opposite-side extract, the two states swap and the ratio inverts (> 1). *Original + decitabine*: DNMT inhibition abolishes the difference (ratio ≈ 1). **b.** Quantification of the L/R MT/DB ratio measured by flow cytometry under the three conditions: Original extracts, Inverted, and Original+Decitabine. The ratio falls below 1 in Original, rises above 1 after inversion (unpaired t-test, p = 0.05), and returns to ≈1 under decitabine. The dashed line marks a ratio of 1 (no L–R difference). Data are shown as mean ± SEM; all values are expressed as the L-treated/R-treated ratio.

Reversibility of this kind, where a cellular state is set by the environment yet can be rewritten when the environment changes, is a hallmark of epigenetic regulation rather than fixed genetic differences. This reasoning, together with our previous findings of L-R epigenetic asymmetries^7^, led us to test whether the bioelectric shift depends on DNA methylation. We therefore assessed whether it persisted under methylation inhibition with decitabine. We treated cells with the DNA methyltransferase inhibitor decitabine during the conditioned culture. Under these treatments, the L–R bioelectric differences were lost (as the L/R MT/DB ratio approaches 1, Figure 7b, *Original extract*+*Decitabine*), confirming that the depolarization bias of L-treated cells is reversible and methylation-dependent.

It is noteworthy that these results could not be reproduced in the xenograft model following contralateral reimplantation (Supplementary Figure 4), an outcome that we attribute primarily to the limitations of the experimental approach. Specifically, transplanting a fragment of a tumor including the stroma infiltration of the original side, could retard/alter the environment signals of the new breast side. A more suitable strategy would likely involve prior ex-vivo tumor dissociation and culture recapitulating the original L–R differences before reimplantation.

Taken together, these results confirm that the L–R bioelectric bias can be reproduced using tissue-derived extracts from different species, reveals for the first time that it is reversible upon altering the extracellular environment (high plasticity) and depends on DNA methylation.

### 3. L-R EPIGENETIC DIFFERENCES IN BREAST TUMORS

#### No L-R differences in global methylomes of Xenograft tumors

We aimed to further explore potential DNA methylation differences in the animal model. To this end, we analyzed the genome-wide DNA methylation profiles of three pairs of xenograft tumors. After performing long-read methylome profiling with the Oxford Nanopore MinIon platform, we found that at the global level, no significant differences in total DNA methylation were observed between L-R tumors (Figure 8a).

**Figure 8.**
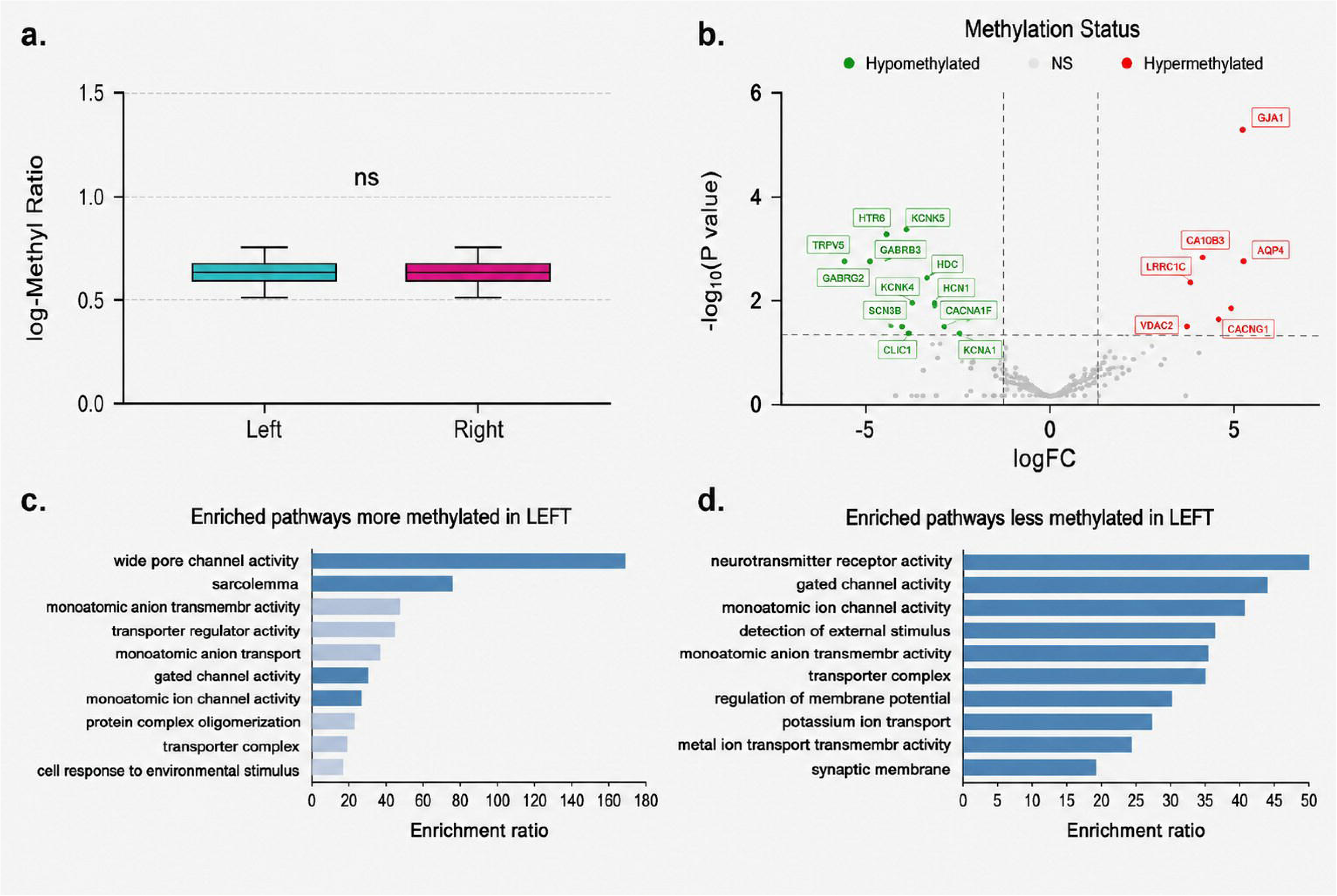
Methylation differences between L-R xenograft tumors. **a.** Global DNA methylation in three L–R pairs of xenograft tumors, determined by Nanopore long-read sequencing and expressed as the log-methylation ratio, did not differ significantly between sides (paired t-test, p > 0.05; ns). **b**. Volcano plot of differential methylation in L versus R tumors (edgeR), showing –log₁₀(P value) against log₂ fold-change. Dashed lines mark the significance thresholds. Coloured points are ICH genes reaching nominal significance — hypomethylated (green) or hypermethylated (red) in L tumors — while grey points are non-significant (NS). *GJA1*, the only gene surviving FDR correction (Table 1), is the most strongly hypermethylated gene in L tumors. **c–d.** Enriched Gene Ontology terms among ICH genes hypermethylated (c) and hypomethylated (d) in L tumors, ranked by enrichment ratio. Analyses performed in R; figure assembled in Adobe Illustrator 2024

**Table 1.**
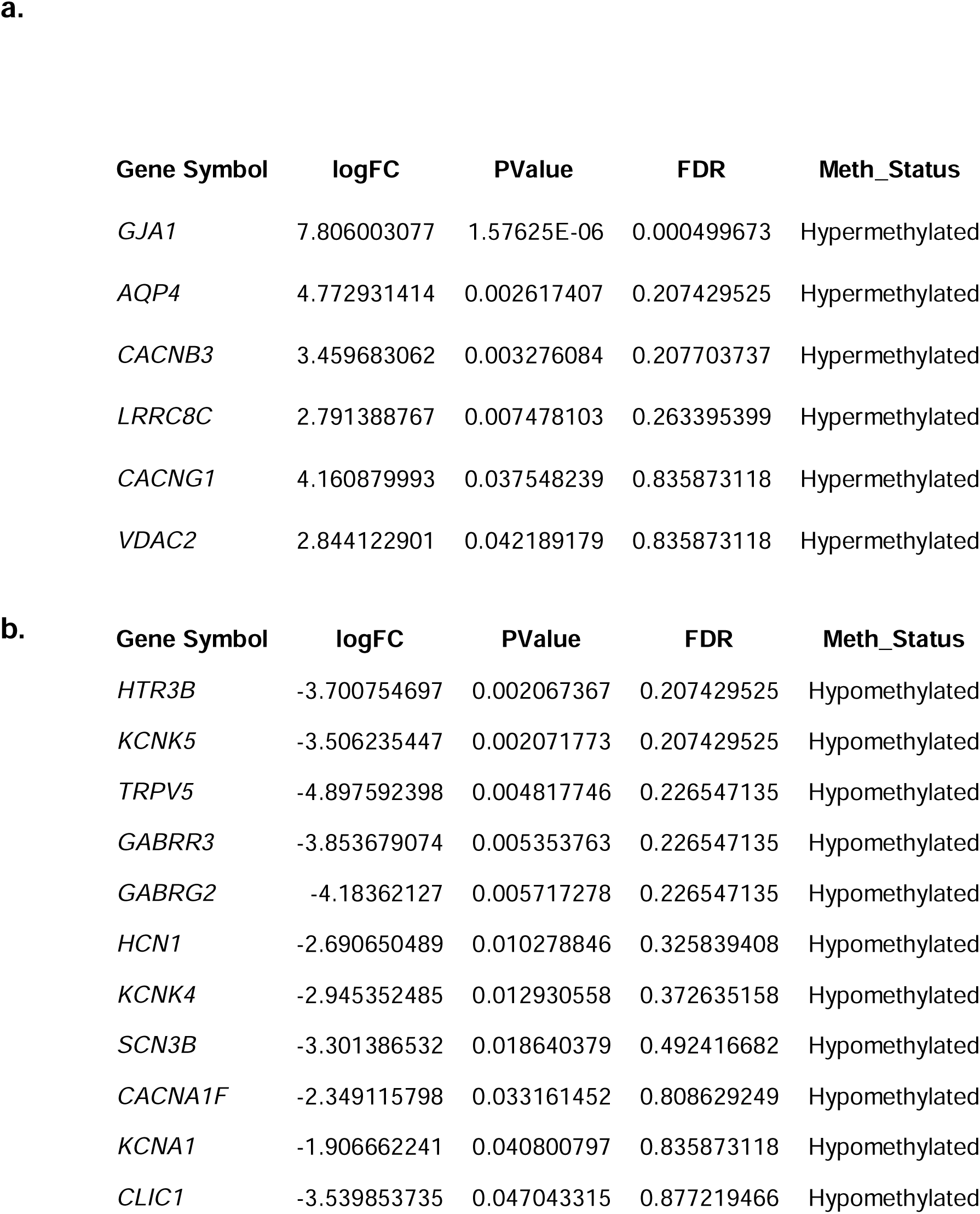
Ion channel (ICH) genes differentially methylated (edgeR, Benjamini-Hochberg FDR correction) comparing L-R xenografted tumors. *GJA1* is the only gene reaching FDR < 0.05. Additional genes shown reached nominal significance (p < 0.05) but did not survive multiple testing correction and are listed as nominally significant candidates. Results are listed as more (a) and less (b) methylated in L tumors compared to R.

#### L-R methylation differences in ion channel (ICH) genes of xenografted tumors

We focused on 228 of the 330 human ICH genes (HGNC classification) in MDA-MB-231 xenografts, building on our prior observation of epigenetic ICH differences in the PyMT mice model^6^. Differential methylation analysis identified *GJA1* as the only gene significantly hypermethylated in L tumors after FDR correction (FDR = 0.00049; Table 1a). Five additional genes — *AQP4, CACNB3, LRRC8C, CACNG1*, and *VDAC2* — were hypermethylated at nominal significance (P < 0.05) but did not survive multiple testing correction, as did eleven hypomethylated candidates in L tumors (*HTR3B, KCNK5, TRPV5, GABRR3, GABRG2, HCN1, KCNK4, SCN3B, CACNA1F, KCNA1*, and *CLIC1*; Figure 8b; Table 1b). Despite the limited sample size, the nominally significant gene set converged on coherent functional categories. Enrichment analysis showed hypermethylated genes were associated with wide-pore and voltage-gated channel activity, while hypomethylated genes mapped to neurotransmitter receptor activity and regulation of membrane potential (Figure 8c–d). *GJA1* encodes Connexin 43, the principal gap junction protein in breast epithelium and a pre-specified candidate based on its role in intercellular electrical coupling — making its confirmation under FDR correction the primary finding of this analysis.

#### Reduced expression of electrical coupling gene signatures in L TCGA tumors

GJA1 hypermethylation in L xenografts pointed directly to gap junction-mediated electrical coupling as the functionally relevant pathway. To test whether this epigenetic signal was accompanied by transcriptional changes in human tumors, we assembled a gap junction and aquaporin gene signature (*AQP1, AQP7, AQP8, AQP11, GJA1, GJB1, GJB2, GJC2, GJD3*, and *GJA9*; restricted to epithelial expressed genes) and interrogated TCGA IDC RNA-seq data via the UCSC Xena Browser. Expression of this signature was significantly reduced in L-sided tumors relative to R (Welch’s corrected t-test, P = 0.04), consistent with increased methylation found in xenografts.

#### *GJA1* and *GJB2* expressions are reduced in L-treated cells and L-Xenograft tumors

To validate these findings *in-vivo*, we quantified the expression of *GJA1* and *GJB2*, (the two most prevalent gap junction genes in breast tissue) by qPCR in xenograft tumors. Both genes showed reduced expression in L-sided tumors compared to R-sided counterparts (one-sample t-test, p < 0.001; Figure 9a-b).

**Figure 9.**
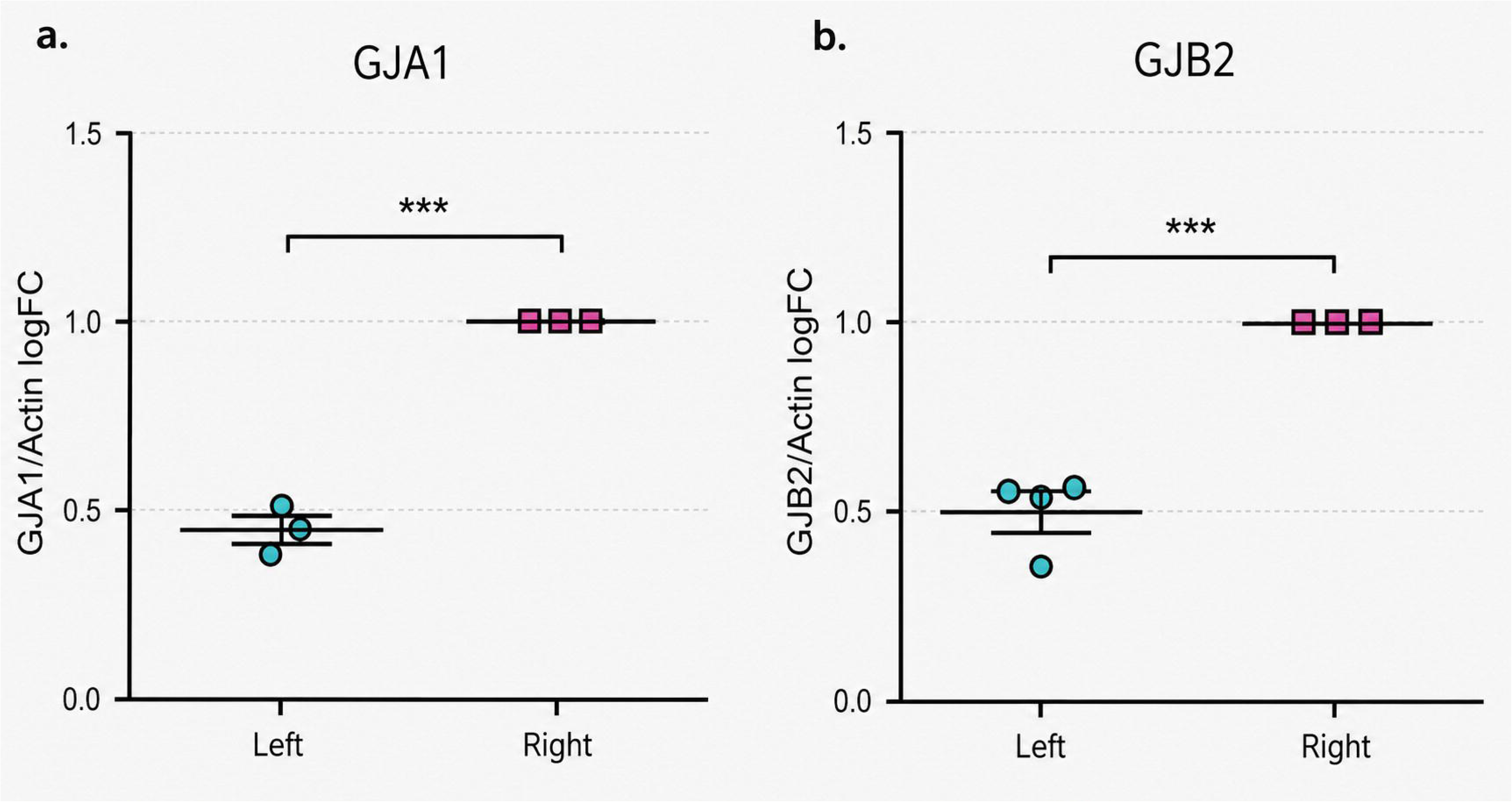
Gene Expression of *GJA1* and *GJB2* in L and R-Sided Xenograft Tumors. Gene expression of *GJA1* and *GJB2* was analyzed in 3–4 pairs of L- and R-sided xenografted tumors. **a-b:** qPCR determination of *GJA1* and *GJB2* expressions show a reduction in L-sided tumors (one-sample t-Test, p < 0,001, *GJA1* n=3 and *GJB2* n=4).

In the L–R conditioned cell culture model, we examined by qPCR *GJA1* expression (as *GJB2* is nearly undetectable in MDA-MB231 cells): L-treated cells showed lower mean expression than R-treated ones, although the difference did not reach significance (unpaired t-test, p = 0.06; Supplementary Figure 5).

Together, these results identify *GJA1* hypermethylation and reduced connexin expression as consistent epigenetic and transcriptional features of L-sided tumors.

#### Proof of principle: Computational model shows gap junction strength governs bistable electrical–epigenetic states

Given the observed L–R differences, we asked whether variations in gap junction strength and DNA methylation are sufficient to generate the two electrical states observed empirically. A computational model developed independently to simulate the evolution of Vmem and methylation in epithelial tissues was applied. We simulated a 10 × 10 cell lattice over 100 ms (with Δt = 0,01 ms), starting from a uniform hyperpolarized state and a global methylation of ICH of 20% (Vmem≈ –60 mV, M = 0.2). Figure 10a maps each cell’s final Vmem in one representative run: nearly all cells return to the hyperpolarized attractor, with only occasional depolarized “islands”. When we repeat the simulation 1000 times (Figure10b), 2 clear outcomes emerge-either the entire tissue remains hyperpolarized or the whole domain flips to depolarization-while mixed states are exceedingly rare. Looking at a single cell across those 1000 runs (Figures 10c-d), its Vmem clusters tightly around two stable states: Vmem≈–55 mV with M≈0.25 (hyperpolarized), or Vmem≈0 mV with M≈1 (depolarized). This bi-stability reflects how small cell-to-cell properties variability (we sampled conductances from a Gaussian distribution) can occasionally tip a cell into depolarization, which then, in certain circumstances, propagates through gap junction coupling.

**Figure 10.**
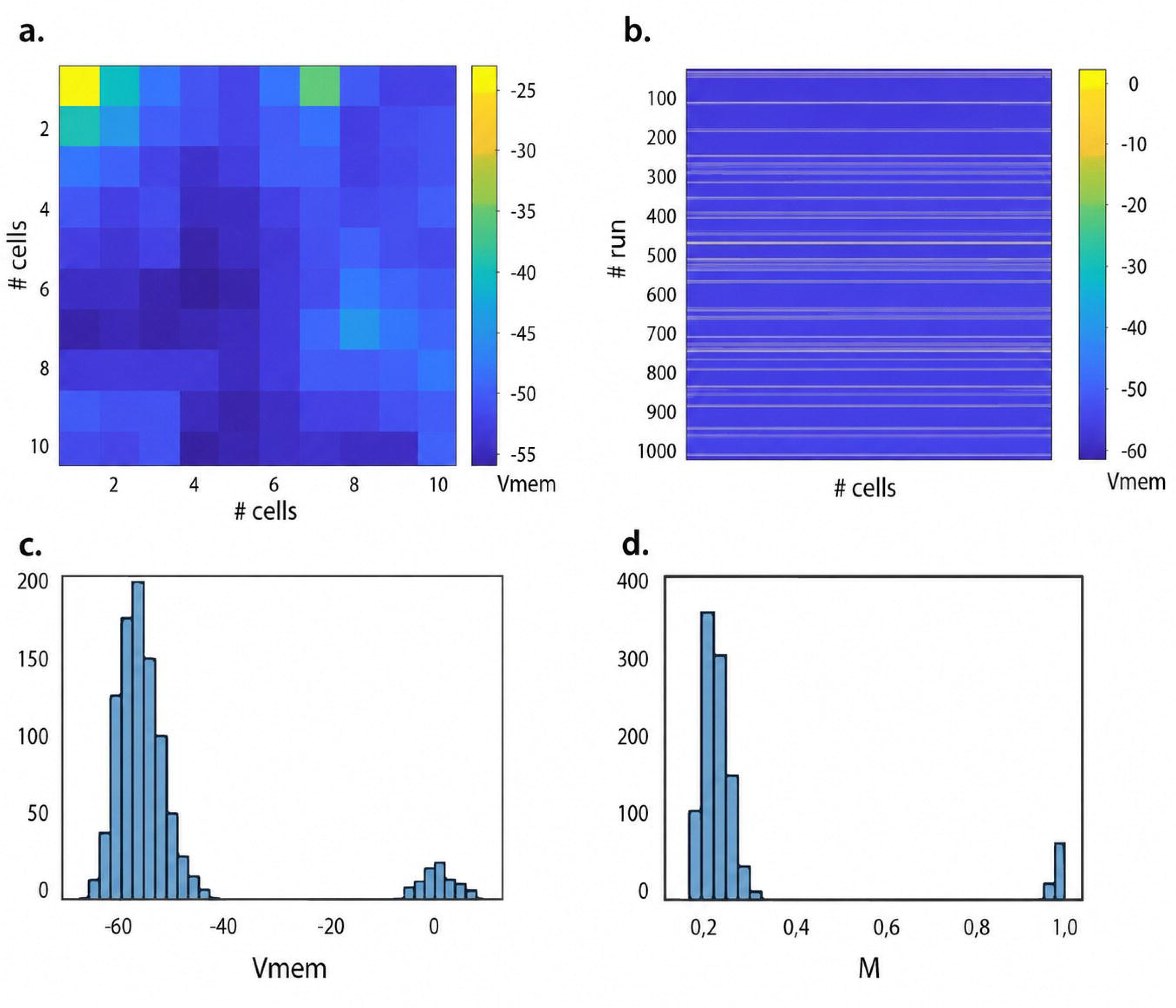
Computer-model outputs demonstrating bistable Vmem and methylation states in a 10×10 tissue. **a**. Final Vmem for each cell in one simulation run. Cells are color-coded by Vmem (blue ≈ –60 mV; yellow ≈ -20 mV). Most remain hyperpolarized, with a solitary depolarized cell visible in the top-left corner (where it is less connected to neighbor cells). **b**. Heatmap of Vmem across all 1000 independent runs (rows) and all 100 cells (columns). Uniform blue bands indicate runs where every cell ended hyperpolarized; uniform orange lines indicate full-domain depolarization—mixed states (dashed lines) are rare, reflecting strong gap-junction coupling. **c-d**. Histograms for a single representative cell over 1000 runs. **c**. Final Vmem clusters tightly around two attractors (≈–55 mV and ≈0 mV). **d.** Corresponding methylation levels bifurcate into two stable values (≈0,25 and ≈1), with negligible occurrences of intermediate states. Figure performed with Adobe Illustrator 2024.

To validate the model’s assumptions, we performed parameter sweeps (Table 2). Decreasing the initial methylation from M = 0.2 toward 0 (hyperpolarization) sharply reduced the chance of global depolarization, whereas moving it toward M = 1 had the opposite effect. Likewise, increasing the depolarizing-channel conductance G0dep from 2.0 to 2.1 nS doubles the frequency of fully depolarized domains; by contrast, changing the polarization-channel conductance G0pol had little effect until it dropped to 0.9 nS, at which point depolarization again became more likely. Interestingly, weakening gap junction coupling (G0max from 1.5 to 1.3 nS)-mimicking the reduced junctional conductance seen in L-cells-increased domain-wide depolarization.

**Table 2:**
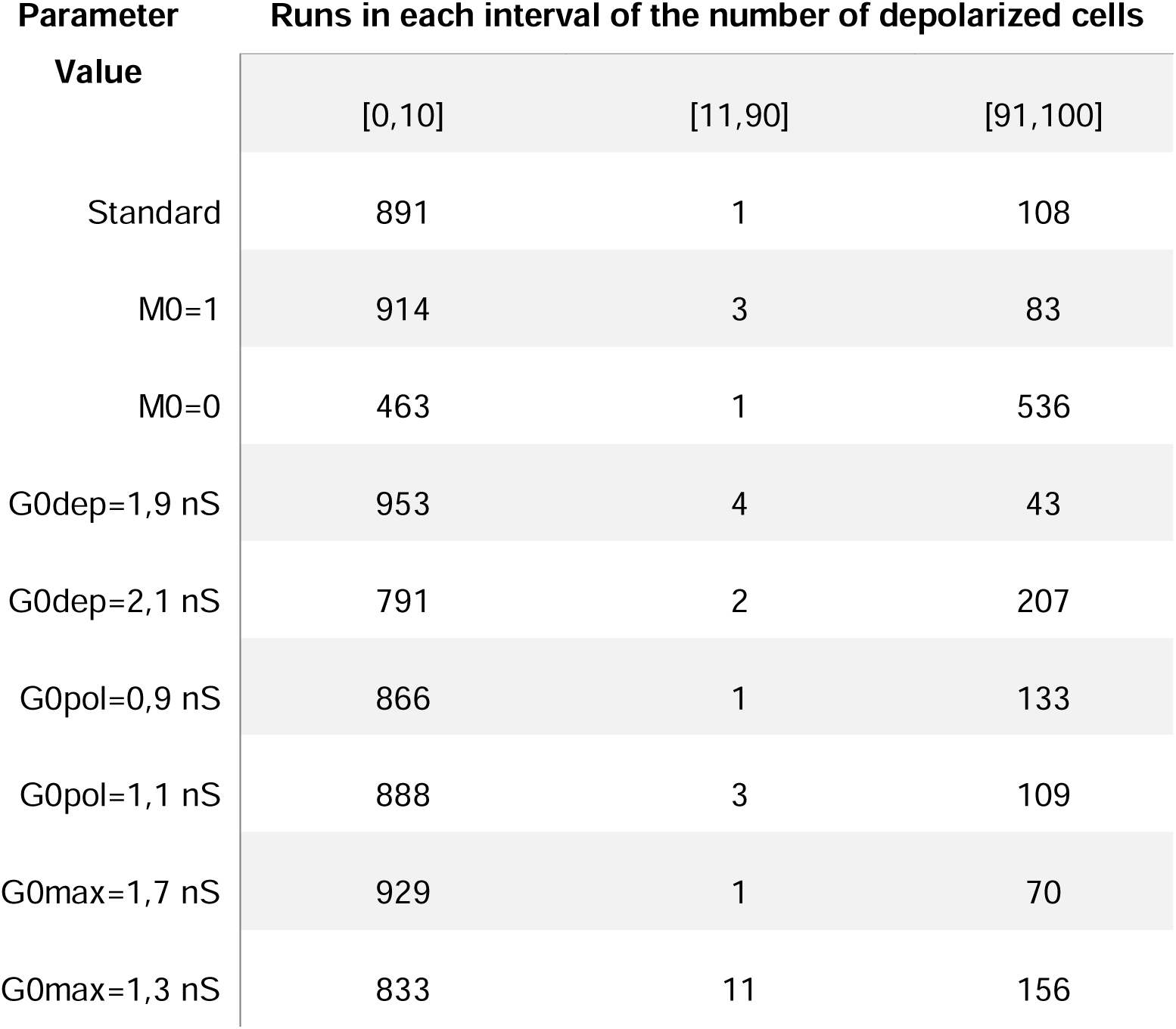
Distribution of the number of depolarized cells at the end of a simulation run, for 1000 runs, with variations of different cells’ bioelectric parameters.

Together, these results support that DNA methylation, by tuning ion-channel conductance, influences the membrane-potential bi-stability that underlie tissue-level L-R asymmetries in breast cancer. They also point to straightforward experimental tests: modulating gap junction conductance pharmacologically or genetically to probe tissue resilience or applying controlled electric fields to assess their impact on methylation-writer expression.

## DISCUSSION

Breast cancer is rarely examined through the lens of L-R anatomy; however, our data suggests that tumors on opposite sides of the body inhabit distinct biological mammary niches and adopt divergent bioelectrical states.

The L-R stromal content analysis of breast tumors identified fibroblast-rich stromal signatures as the principal contributors to the observed laterality differences, showing a modest but remarkably consistent enrichment in R-sided tumors across independent datasets and experimental models. CAFs are a core component of the TME and secrete matrix-modifying enzymes (e.g. matrix metalloproteinases) and extracellular matrix (ECM) components like collagen and fibronectin^29^. This increases the ECM density, resulting in a stiffer matrix that generates compressive forces on tumor cells. As shown by Labernadie et al.^30^, CAFs are able to generate mechanical forces that are transmitted to cancer cells and higher CAF number amplifies the effect. Considering the L-R CAF difference found in our results, we speculate that R-sided tumors may experience stronger mechanical forces and/or be conditioned by distinct soluble factors generated within the TME.

An increased iCAF burden in R-sided tumors could also suggest a TME skewed toward chronic inflammation and tissue remodeling. This pro-inflammatory context could influence immune cell recruitment and activation, potentially affecting the response to immunotherapies. Notably, recent studies have shown that CAF subtypes, including iCAFs, can modulate the efficacy of immune checkpoint inhibitors targeting the PD-1/PD-L1 axis^31^.

Here we propose functional impacts of the TME asymmetries as a consistent hyperpolarization in the R-sided tumors, as compared to the L-counterparts. How stromal composition shapes tumor cell Vmem remains to be experimentally explored. At tissue level, considering that CAF abundance can affect TME density, it is noteworthy that others have recently found that Vmem can change as a response to *mechanical pressure*. Specifically, the authors tested whether changes in biomass density were correlated with changes in Vmem and found that cultured cells became increasingly hyperpolarized with increasing cell number^32^. Thus, the L-R Vmem differences found in our animal models, may reflect a causal link between stromal density and Vmem. The reproducibility of this effect in cell cultures treated with tissue extracts-where intact cells are absent, but their components remain-suggests the involvement of a diffusible molecule rather than a mechanical force. We propose that both factors contribute: variations in mechanical forces combined with differences in diffusible factors. Further studies analyzing the composition of the tissue extracts will help clarify these hypotheses.

Limitation of our work are the small sample in vivo sample size and the modest magnitude of the stromal and CAF asymmetries detected in human cohorts. We therefore interpret them not as a dominant driver of tumor behavior but as one previously unrecognized layer of L-R heterogeneity whose reproducibility is the relevant claim rather than effect size.

The reversibility of the bioelectric shift upon extract inversion, and its abolition by decitabine, provides an initial causal and mechanistic handle, but a full dissection of the ICH and gap-junction circuitry will be required to establish causality The finding aligns with our initial, serendipitous observation in breast cancer patients and reinforces the idea that epigenetic asymmetries can arise even in controlled animal models. In the context of a differential TME, such epigenetic alterations are likely to exert distinct functional impacts on the tumors. The modification of the tumor epigenetic program is a hallmark of cancer^33^ that initiates and propagates tumorigenesis. Altered DNA methylation, histone modifications and ncRNAs expression are a feature of cancer cells. Given the reversible nature of epigenetic modifications, the ability to restore the cancer epigenome through the inhibition of the epigenetic modifiers is a promising therapy for cancer treatment. Since 2 decades ago epigenetics has been suggested as the mechanism linking the environment to changes in gene expression that do not require changes in DNA sequence^34^. It is well known that environmental influences can establish, erase, or reshape epigenetic marks (e.g. DNA methylation, histone modification) and in consequence change gene expression. In this context, our findings suggest a potential L-R differential epigenetic profile that influences tumor cell bioelectric states.

Among the biological processes affected by the involved differential methylated genes, the main asymmetry was found in genes involved in cell electrical coupling. The reduced expression of connexins *GJA1* (Cx43) and *GJB2* (Cx26) in L-sided xenografts supports a functional silencing of electrical coupling. Because gap junctions distribute small ions and second messengers^35^, their down-regulation would effectively decouple L tumors from tissue-level voltage averaging, allowing depolarization to persist in the face of neighboring hyperpolarized cells. In contrast, R tumors apparently retained connexin expression and a Vmem close to that of normal mammary tissue, consistent with the idea that intact cell communication confers electrical homeostasis. Fifty years ago, tumor cells were found to reduce electrical coupling, leading to the hypothesis that loss of direct intercellular communication is associated with cancer progression, a phenomenon later attributed to gap junctions^36^. Connexins regulate not only cell-cell communication but are also implicated in proliferation, apoptosis, chemoresistance, migration and invasion^40^. Connexin-43 (Cx43) is the most characterized gap junction protein and mainly expressed in breast epithelium, primarily involved in intercellular communication between adjacent cells and with the ECM^37^.

The computational model, developed here independently of the empirical work, captures this logic: introducing small variations in depolarizing conductance or methylation, pushes toward one of two attractors—global hyperpolarization with low methylation or widespread depolarization with high methylation—and the outcome is strongly modulated by gap junction coupling strength.

From a clinical standpoint, lateralized biology raises the possibility — not yet established — that prognostic markers and therapeutic vulnerabilities could differ between breasts. Our previous work showed that R tumors proliferate less rapidly^11^, and the present associations are consistent with a contribution from iCAF dominance and preserved electrical coupling, features reported elsewhere to restrain tumor growth or prime immune responses; however, our data do not establish this link causally. The converse combination in L tumors — depolarization, connexin silencing, and dCAF enrichment — aligns with hallmarks associated with proliferation and invasion, though again the association is correlative. Should these lateralized CAF and bioelectric differences be confirmed in larger, clinically annotated cohorts, they could in principle offer a new angle for patient stratification or therapeutic targeting. We emphasize that this remains a hypothesis to be tested rather than a clinical recommendation.

Speculatively, bioelectric modulation and epigenetic editing could represent actionable levers worth exploring: agents that reopen gap junctions or hyperpolarize Vmem might, in principle, shift L tumors toward an R-like state. A next interesting step is testing whether such interventions alter tumor behavior.

Beyond breast cancer, our findings may prompt a reevaluation of paired organs in oncogenesis, consistent with a broader view in which bioelectric and epigenetic profiles are shaped in part by body location — a possibility that will require direct testing in other paired-organ systems.

Whereas earlier work established that the mammary gland is intrinsically lateralized at the transcriptomic level, with prognostic relevance^38^, our results extend this asymmetry beyond the tumor itself to the microenvironment, linking it to coordinated stromal, bioelectric, and epigenetic differences. Altogether, building on prior evidence, our findings reveal that breast tumor laterality extends to the microenvironment, integrating stromal, bioelectrical, and epigenetic dimensions, and underscoring how spatial context within the body can shape tumor biology.

## METHODS

### Extracts from human and bovine mammary tissue

All extracts used in this study were derived exclusively from healthy, non-tumoral mammary tissue of two species — human and bovine — and were applied only to cultured MDA-MB-231 breast cancer cells. No tumor-derived or tumor-conditioned extracts were used. L and R-sided tissue was processed separately throughout, so that every extract retains its side of origin. Four healthy L-R human breast glands were obtained from plastic surgeries, provided by Dr. Cataneo from the Clinic of Plastic Surgery of Mendoza, after patients signed an informed consent previously approved by the Ethics Committee of the Medical School of the National University of Cuyo, according to the guidelines of the Declaration of Helsinki (Protocol code: 15856/2016, date of approval: 3 December 2019). Tissues were first disaggregated with a scalpel and the pieces were suspended in 25 mL of DMEM medium with Penicillin/Streptomycin 1% and incubated in a shaker for 24 h at 37 °C. Next, samples were centrifuged to remove solid fat, and the remaining suspension was filtered with cell strainers of first, 100 μm and afterwards 40 μm, to eliminate residual tissue parts. The obtained liquid phase extracts were L-R labeled and stored for further experiments at − 20 °C. For further experiments, a pool of the 4 extracts was mixed to condition cells in culture.

L-R bovine udder samples were obtained from 2 nulliparous heifers immediately after slaughter at the Frigorífico María del Carmen abattoir (https://frigorificomariadelcarmen.com.ar/home/, certified by SENASA). From the collected tissues, the udder was separated, sectioned, and stored in sterile containers at -20°C until processing. Tissues were mechanically disaggregated using surgical scissors or an Ultraturrax (high-speed rotor) until reaching a weight of 22 g. The material was then transferred to a 50 ml Falcon tube and mixed with 25 ml of DMEM/F-12 culture medium supplemented with 100 U/ml penicillin and 100 µg/ml streptomycin. The resulting extract was incubated at 37°C on a horizontal shaker for 24 hours. Subsequently, the solid and liquid components were separated using a cell strainer and cellulose acetate filters of 5 µm, 0.45 µm, and 0.22 µm pore sizes (the final filter step performed under a biosafety cabinet to ensure sterility). The resulting liquid-phase extract was stored at -20°C until use.

### Conditioned cell culture

Human breast cancer cell line MDA-MB-231 (ATCC, RRID:CVCL_0062) was obtained from the Tissue and cell bank from CIQUIBIC Institute (UNC-CONICET), Cordoba, Argentina, and passages up to 15 were used for this work. In general, cells were cultured in DMEM medium (Gibco by Life Technologies, Grand Island, NY, USA, # 112800-058) supplemented with 10% fetal bovine serum (Internegocios S.A, Mercedes, BA, Argentina), 100 U/mL of penicillin and 100 μg/mL streptomycin (Gibco by Life Technologies, Grand Island, NY, USA, #1796440), at 37 °C in a humidified atmosphere containing 5% CO2. For the extract-conditioned cultures, fetal bovine serum was reduced to 1%. MDA-MB-231 were conditioned with a cocktail of different proportions depending on the tissue origin: cocktails from human tissue source consisted of 49% DMEM with Penicillin/Streptomycin, 1% Fetal Bovine Serum and 50% L/R liquid-phase extract; while cocktails from bovine source had 69% DMEM with Penicillin/Streptomycin, 1% Fetal Bovine Serum and 30% L/R liquid-phase extract.

### Xenograft and PDX tumors

All animal procedures were reviewed and approved by the CICUAL committee of the National University of Cuyo, Mendoza, Argentina. 8-10 weeks of age NodScidGamma (NSG) female mice (Jackson Laboratory, Bar Harbor, ME, USA) were housed together in the same cage in an on-site housing facility, with access to standard food, water, and a 12/12 light cycle. For the xenograft assay, 10^6^ MDA-MB-231 cells were inoculated in the 4^th^ L-R mammary glands of each NSG mice. When the tumors developed and reached 1000 mm^3^ size, the mice were euthanized using CO_2_, and paired L-R breast tumors were excised. Tumors were fractionated into five 1.5-2 mm slides for measuring Vmem by confocal microscopy and for DNA/RNA extraction. The patient-derived xenograft (PDX) BC-AR474 model was kindly provided by Dr. Claudia Lanari (IBYME-CONICET, BsAs, Argentina)^41^. The PDX tumor was grown and maintained in NSG mice by fractionating and retransplanting 2mm^3^ tumor portions into the L-R 4th mammary glands. When the PDX tumor reached 1000 mm^3^, the mice were sacrificed with CO₂.

### Immunofluorescence

Paired mammary tumors L-R were fixed with 4% formalin 24 h at 4 °C (ANOO6739PI, Anedra, BsAs, Argentina), and then incubated with increased 10-20-30% sucrose (Sigma Aldrich) in PBS for 24 h at 4 °C each step. Subsequently, the tumors were mounted in OCT compound (Cellpath 050226) and maintained at -20 °C until use. OCT-prefixed PDX/xenograft tumors were sectioned at 20 µm wide using a microtome at -22°C and mounted on positively charged slides at room temperature. The tissue was permeabilized with 3% Triton 20 in 100% methanol, blocked with 5% fetal bovine serum in PBS, and incubated overnight with a 1/100 dilution of the primary antibody against alpha-smooth muscle actin (α-SMA, DACO M0851) in blocking solution. Subsequently, it was incubated for 3 h with 1/750 dilution of secondary antibody Goat Anti-Mouse IgG H&L (Alexa Fluor® 647, ab150115) in blocking solution, and the nucleus was stained for 10 minutes with Hoechst 33342 (Thermo Scientific 62249) at a 1/1000 dilution in PBS. Five/ten images of each tumor were captured with a 20X objective and examined by fluorescence confocal microscopy (Olympus FV1000-EVA® confocal microscope). For the analysis, the total α-SMA signal was normalized to total Hoechst signal or to total nuclei, for each image capture, using Image J software analysis (ImageJ 1.53t/Java 1.8.0_322, National Institutes of Health, USA, RRID: SCR_003070). Paired values were analyzed using a two-tailed paired Student’s t-test.

### Flow Cytometry of tumor single-cell suspensions

Single-cell suspensions were prepared from freshly excised tumor tissues. Briefly, tumors were minced into small fragments and digested for 40 min at 37 °C in PBS containing collagenase D (1 mg/mL; Roche) and DNase I (100 µg/mL; Roche). The digested material was filtered through a 70 µm cell strainer, washed with PBS containing 2% fetal bovine serum (FBS), and centrifuged at 400 × g for 5 min. Red blood cells were lysed using ACK buffer when necessary. Cells were subsequently stained for 20 min at 4 °C with fluorochrome-conjugated antibodies against the following markers: CD45 (clone 30-F11) to identify total leukocytes (immune cells); CD11b (clone M1/70) to define myeloid cells; CD11c (clone HL3) and F4/80 (clone BM8) to discriminate dendritic cells within the CD11b⁺ gate; CD140a (clone APA5) and CD31 (clone MEC13.3) to distinguish fibroblasts (CD140a⁺ CD31⁻) and endothelial cells (CD31⁺) within the non-immune stromal (CD45⁻) fraction. Data acquisition was performed on a FACS Arias flow cytometer (BD Biosciences), and data was analyzed with FlowJo v10.7.1 software. Fluorescence parameters were displayed on biexponential scales, while forward and side scatter were represented linearly. The gating hierarchy is illustrated in Figure 5a.

### Vmem analysis

#### Vmem Measured by Flow Cytometry

As previously described by us^6^, MDA-MB-231 cells were plated and conditioned with L-R extracts for 5 days, trypsinized and then incubated for 30 min with 2 μM DiBAC4(3) (Bis-1,3-Dibutylbarbituric AcidTrimethine Oxonol, an Alexa-fluorescent probe negatively charged) (Invitrogen by Thermofisher Scientific, Cat. No. B438) and 1 μM MitoTrackerRed (an APC-fluorescent probe positively charged) at 37 °C and 5% CO2. Afterwards, cell fluorescence was measured by flow cytometry (FACSARIA-III, BD-Biosciences®) with a BP 530/30 emission filter. Results were analyzed using FlowJo v X.0.7® software (RRID:SCR_008520). To provoke maximum depolarization, cells were first treated with 65 mM KCl for 5 min at 37 °C and afterwards incubated for 30 min with the fluorescent probe DiBAC4(3) and MTRed as described above. For Vmem comparisons of cultured cells, 10.000 events per sample were acquired by flow cytometry in each biological replicate. L-R differences are calculated as the L Mitotracker/Dibac ratio normalized to the R-ratio.

#### Vmem Measured by Confocal Microscopy

Thin slices of the paired L-R 4th gland tumors were cut and placed in Petri dishes with DMEM FluoroBrite (Gibco, #A1896701, Life Technologies, Carlsbad, CA, USA) plus 2% FBS. For measuring Vmem, 2–3 slices per tumor were used, incubating 15 min at 37 °C with media containing DB 2 mM and MT 500 nM in Fluorobrite DMEM. One slice was used to measure autofluorescence of the cells. Images were captured using a Leica Stellaris Sp8 confocal microscope at 10X magnification, at 37 °C and 3–5 images were taken per tumor, with 3 z-slices per image. As control for normalized measurements, cells were treated with a high concentrated KCl (65 mM) solution as depolarizing agent, for 15 min at 37 °C. Afterwards, slices were incubated with Fluorobrite DMEM containing DB (2 mM) and MT (500 nM). The images were processed using Image J software (ImageJ 1.53t/Java 1.8.0_322, National Institutes of Health, USA, RRID: SCR_003070). For Vmem comparisons in mouse xenograft tumors, confocal Z-stacks from three paired tumors were collapsed to summed-intensity projections. Mean fluorescence intensities were extracted using ImageJ v1.54, and inter-side comparisons were made using a two-tailed One-sample t-test against 1. L-R differences are calculated as the L Mitotracker/Dibac ratio normalized to the R-ratio.

### DNA extraction

Cells were collected and tissues were homogenized in PBS using an Ultra turrax homogenizer, following centrifugation, the pellet (cells or tissue) was suspended and washed 1 time with Tris–EDTA (T_10_ E_10_ Buffer), suspended in cetyltrimethylammonium bromide (CTAB) solution (2 g/l CTAB Sigma-Aldrich®, 100 mM Tris/HCl, 20 mM EDTA and 2% 2-mercaptoethanol) and incubated at 60°C for 1 h for cells or overnight for tissue. Then, chloroform-isoamyl alcohol solution (24:1) was added and the sample was centrifuged. The aqueous phase was collected into a new tube and mixed with 3 volumes of ice-cold 100% ethanol. Precipitated DNA was dissolved in T_10_ E_0,1_ Buffer. Further the samples were cleaned and purified using QIAamp DNA Micro Kit (Qiagen), according to manufacturer’s instructions.

### RNA extraction and qPCR gene expression analysis

Total RNA from cells and xenograft tumors were extracted using a cell disruptor (Pro200 Proscientific), and TriPure Isolation Reagent (11667165001, Roche), following the manufacturer’s protocol. RNA concentration and purity were determined using a NanoDrop spectrophotometer (Labocon). One microgram of RNA was reverse-transcribed to cDNA using M-MLV Reverse Transcriptase (INBIO Highway). cDNA was diluted to 4 ng/μL and stored at -20 °C. Real-time PCR was conducted with 16 ng cDNA per reaction using the Master qPCR SYBR Green Kit (INBIO Highway). Primer sequences used were: *GJA1* forward: 5’-TGAGTGCCTGAACTTGCCTT-3’, *GJA1* reverse: 5’-GCCTGGGCACCACTCTTTT-3’; GJB2 forward: 5’-CTCCCGACGCAGAGCAAA-3’, GJB2 reverse: 5’-TCATCTCCCCACACCTCCTT-3’; and ACTB (β-actin) forward: 5’-TGACGTGGACATCCGCAAAG-3’, ACTB reverse: 5’-CTGGAAGGTGGACAGCGAGG-3’. The PCR reaction was performed in Aria Mx-Real time PCR system/Agylent technologies®, and the analysis was performed using AriaMx Software version 2.0 (Agilent, Santa Clara, CA, USA). Relative expression normalization of genes of interest was carried out using β-actin gene expression as endogenous reference control by the ΔΔCq method. Technical triplicates were averaged per sample. Expression ratios between L and R tumors for *GJA1* and *GJB2* were evaluated using a one-sample two-tailed t-Test against a theoretical ratio of 1.

### Methylome analysis

#### Library preparation and sequencing

∼200 ng DNA samples from xenograft tumors conditions (L and R, 3 paired tumors) were used as input for Nanopore long-read sequencing. Duplicate libraries were prepared to obtain higher coverage. The libraries were prepared using the Rapid Barcoding Kit from Oxford Nanopore Technologies, following the manufacturer’s instructions. Briefly, this system employs transposome complexes that cleave the DNA and add barcodes to each sample. The samples are then pooled, and the library is purified using magnetic beads. Finally, sequencing-specific adapters are added. Sequencing was performed on a MinION Mk1C device for 48 hours at 37°C, with no real-time basecalling nor alignment, generating POD5 files for subsequent analysis.

#### Raw data processing, global methylation and differential methylated region analysis

POD5 files were processed for base calling using Dorado basecaller v0.9.1, including the identification of 5-methylcytosine and 5-hydroxymethylcytosine using the high-accuracy model (HAC). The resulting .bam files were aligned to the reference genome hg38, obtained from the UCSC Genome Browser (https://genome.ucsc.edu/), using Dorado aligner (minimap2 v2.27). Subsequently, individual .bam files were sorted and indexed, then the duplicated .bam files from each sequencing were accordingly merged, re-sorted and re-indexed using SamTools v1.13. Additionally, for further analyses, bedmethyl files containing the genomic coordinates of all modified cytosines were obtained using ModKit v0.4.4. For the bioinformatics analysis, the NanoMethViz v2.8.1 package was initially used to generate a Tabix files containing the chromosomal coordinates of each detected cytosine modification and its Log-Likelihood Ratio (LLR), which indicates the probability of cytosine modification. For global methylation analysis the methy_to_bsseq() and bsseq_to_log_methy_ratio() functions were used to convert the LLR of each genomic position into a logarithmic methylation ratio (LMR). Prior to statistical analysis and plot generation, the log-methylation ratios were transformed into values ranging from 0 (unmethylated) to 1 (methylated) using a sigmoidal function: 1/1+e ^-LMR^. For differential methylated region analysis (DMR) the methy_to_bsseq() and bsseq_to_edger() functions were used to convert the LLR of each genomic position into an edgeR methylation matrix. Coverage estimation, normalization, experimental design modelling, verification of experimental design, estimation dispersion and Likelihood test were performed using edgeR package v4.0.16. Plots were modelled using ggplot2 v3.5.2 and ggpubr v0.6.0 packages. Data and tables were formatted and modelled using reshape2 v1.4.4, tidyr v1.3.1, dplyr v1.1.4 and stringR v1.5.1 packages. Statistical analyses were performed with Rbase v4.3.2. All R scripts used for the analysis are available in the ‘Code Availability’ section.

### Bioinformatic analysis

Clinical and RNA-seq data from 1096 breast primary tumors were acquired from TCGA dataset in Genomic Data Commons repository (https://portal.gdc.cancer.gov/) using TCGAbiolinks R library. Samples were filtered by tumor type, yielding a final cohort of 784 invasive ductal carcinoma (IDC). These were further filtered to retain only those with a CPE ≤ 70%, obtained using the tumor.purity function from the TCGAbiolinks v2.30.4 package, resulting in a cohort of 276 samples. The cohorts of 784 (424 L – 360 R) and 276 samples (153 L – 123 R) were then classified according to anatomical site (L–R). To assess the presence of stromal and immune cells, the xCell v1.1.0 R package was used, which assigns an immune score and stromal score based either on predefined gene signatures from 64 stromal and immune cell types or immune score signature and stromal signature only (gene list details in Supp Data1). Plots were modelled using ggplot2 v3.5.2 and ggpubr v0.6.0 packages. Data and tables were formatted and modelled using reshape2 v1.4.4, tidyr v1.3.1, dplyr v1.1.4 and stringR v1.5.1 packages. Normalization of RNAseq count by gene-length method and outliers’ analysis were performed with EDAseq package v2.36. Gene annotations were performed via biomaRt v2.58.2 package. Statistical analyses were performed with Rbase v4.3.2. All R scripts used for this analysis are available in the ‘Code Availability’ section.

For nuclear segmentation and classification analysis in H&E histological slides, 10 L and 10 R IDC samples from the cohort of 784 analyzed by Xcell were selected. The corresponding diagnostic slides were retrieved from the Imaging Data Commons Portal (https://portal.imaging.datacommons.cancer.gov) in whole slide image (WSI) format (.svs). For each WSI, a region of interest (ROI) of 2000 µm × 2000 µm was defined using QuPath v0.6.0 (https://qupath.github.io/), prioritizing areas with high nuclear density and based on predefined technical quality criteria (e.g., staining consistency, absence of artifacts) while blinded to tumor laterality and outcomes. Each ROI was then subdivided into non-overlapping tiles of 512 px (≈260 tiles per ROI) using the OpenSlide v1.4.2 Python library. All tiles were processed with the PyTorch v2.2.0 implementation of HoVer-Net, a deep learning framework for nuclear segmentation and classification of H&E-stained images. HoVer-Net was applied using a pretrained model based on the PanNuke dataset^39^, which includes ∼790,000 manually annotated nuclei across 19 tissue types, including breast carcinoma. This model classifies nuclei into five categories: Neoplastic (tumor), Connective (fibroblast/stroma), Inflammatory, Non-neoplastic epithelial (normal), and Dead (necrotic/apoptotic). Output from HoVer-Net consisted of comma-separated value (.csv) files containing X–Y nuclear coordinates and the predicted nuclear type for each tile. Data from all tiles per sample were merged, and cell type frequencies were computed per sample and per anatomical side (L vs R). Comparisons between L and R cohorts were performed either by stromal-to-tumor ratio (Connective/Neoplastic) or by stromal cell abundance relative to stromal and epithelial cells (Connective cells/(Neoplastic cells + Connective cells). Non-neoplastic epithelial cells were excluded from this calculation because they represent residual normal breast tissue rather than components of the tumor microenvironment. Statistical analyses were performed with Rbase v4.3.2. All R scripts used for this analysis are available in the ‘Code Availability’ section.

### Computational modeling

A simple computer model was developed and run to describe the L-R asymmetries found in breast cancer. It is a cellular automaton, an adaptation of previously published models, as in reference^40^.

Cells can exchange ions with the extracellular medium by general depolarization and polarization ICHs, and the intercellular communication is described by a universal gap junction. The activity of each of these components depends on the membrane voltage, as will be shown in the following equations. The electrical potential (relative to the cell exterior) of cell *i*, V_i_, changes due to ionic currents (considering only the movement of positive ions). Its variation with time is determined by the ordinary differential equation,

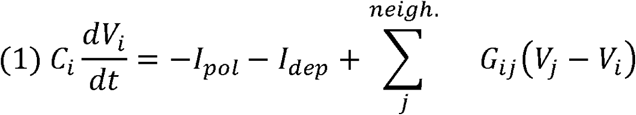

where C_i_ is the cell membrane capacitance, I_pol_ the current through polarization ICHs, I_dep_ the current by depolarization ICHs, and G_ij_ the conductance of ion transfer through gap junctions between contiguous cells *i* and *j*. The sum is done over all the neighbor cells, if present, considering a von Neumann neighborhood. The currents and conductances on the expression depend on the membrane electric potential as follows,

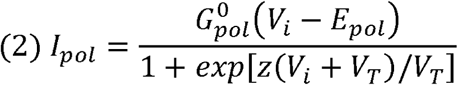

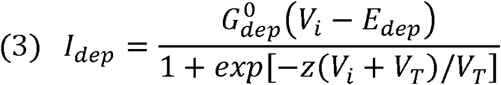

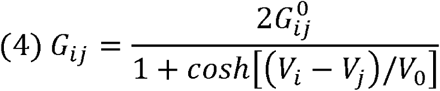

with

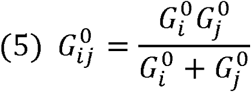

and

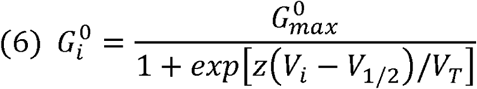

where E_pol_ (E_dep_) is the polarization (depolarization) membrane electric potential, V_T_ the threshold potential, *z* the channel gating charge, G^0^_pol_ (G^0^_dep_) the polarization (depolarization) ICHs conductance, V_1/2_ the value of electrical potential that reduces G_i_^0^ by a factor 2, and V_0_ adjusts the gap junction conductance function width^39^. The model parameters were determined, based on experimental results, and presented in reference^39^ for a generic non-excitable cell; the values used in the current work are given in Table 3. As previously shown, G_ij_ describes the gap junction ion conductance between adjacent cells, representing the results of the serial association of cells *i* and *j* connexons conductances (G_i_^0^ and Gj^0^).

**Table 3:**
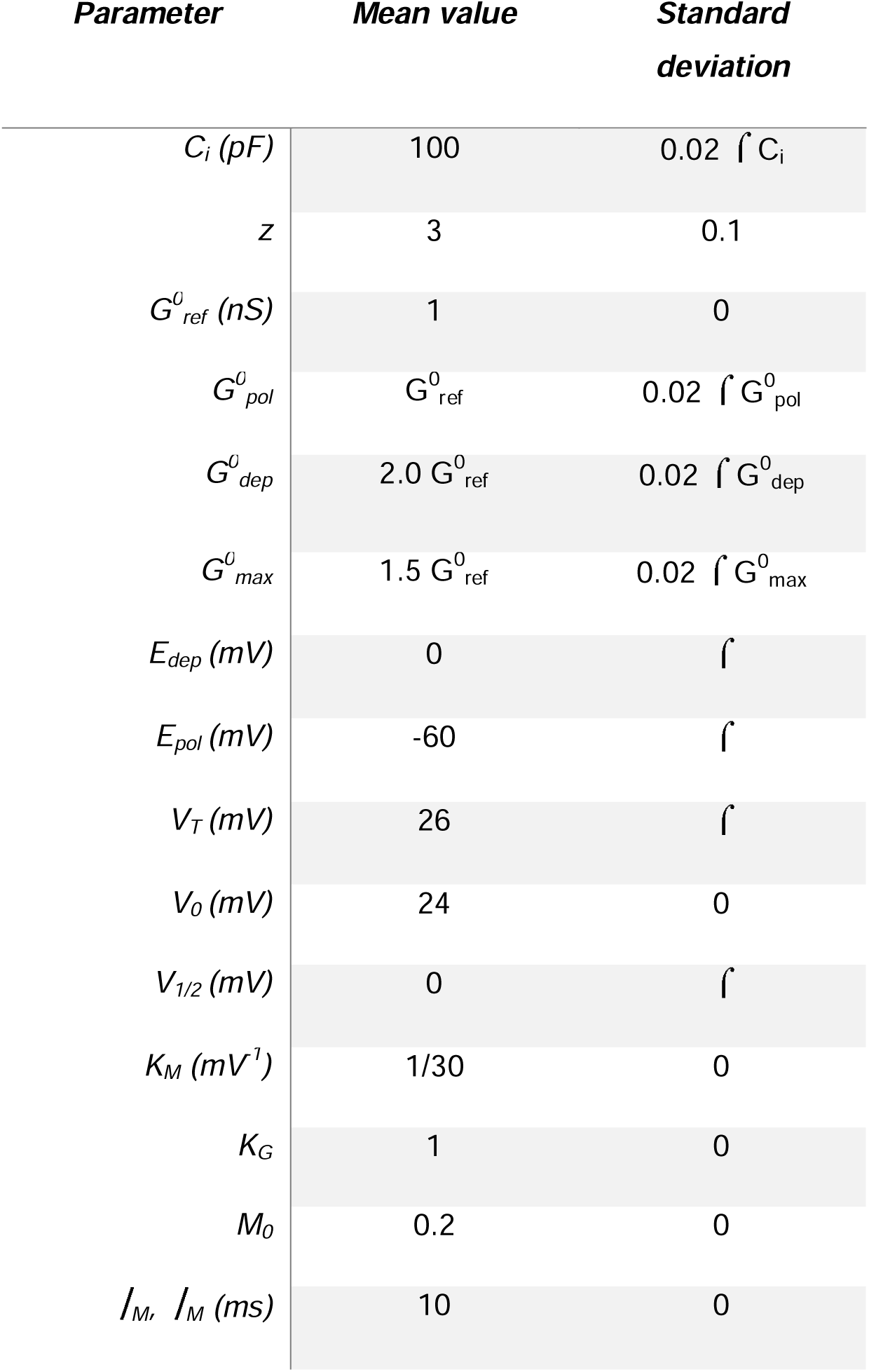
Values of the different cells’ bioelectric parameters used on the present model. ⌠=3 is used as the standard deviation on gaussian distributed values around the mean value (stochastic model).

| **Parameter** | **Mean value** | **Standard deviation** |
| --- | --- | --- |
| $C_i$ (pF) | 100 | $0.02 \hat{\sigma} C_i$ |
| $z$ | 3 | 0.1 |
| $G_{ref}^0$ (nS) | 1 | 0 |
| $G_{pol}^0$ | $G_{ref}^0$ | $0.02 \hat{\sigma} G_{pol}^0$ |
| $G_{dep}^0$ | $2.0 G_{ref}^0$ | $0.02 \hat{\sigma} G_{dep}^0$ |
| $G_{max}^0$ | $1.5 G_{ref}^0$ | $0.02 \hat{\sigma} G_{max}^0$ |
| $E_{dep}$ (mV) | 0 | $\hat{\sigma}$ |
| $E_{pol}$ (mV) | -60 | $\hat{\sigma}$ |
| $V_T$ (mV) | 26 | $\hat{\sigma}$ |
| $V_0$ (mV) | 24 | 0 |
| $V_{1/2}$ (mV) | 0 | $\hat{\sigma}$ |
| $K_M$ (mV<sup>-1</sup>) | 1/30 | 0 |
| $K_G$ | 1 | 0 |
| $M_0$ | 0.2 | 0 |
| $I_M, I_M$ (ms) | 10 | 0 |

Cells are not exactly equal, and in this model, the main individual cell bioelectric parameters magnitude follows a Gaussian distribution. In Table 3 the second column presents the parameter mean value and the third column the respective standard deviation.

The effect of an increase of DNA methylation observed in the 6 ICH-signature described by us previously (the 6-ICH signature^7^) of depolarized cells, is introduced in the following equation (Hill equation, with Hill coefficient n=2),

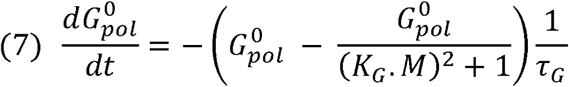

where the conductance of the polarization channels (G^0^_pol_) decreases with an increase on the level of ICH methylation. This is represented by the variable *M*, with a value between 0 and 1. The parameter K_G_ gives the inverse of the value of methylation that decreases G^0^_pol_ change rate by half of the reference value G^0^_ref_. ⎮_G_ gives the time scale of change in G^0^_pol_. An increase of *M* brings a decrease of the stable value of G^0^_pol_ and then to a lower polarization current and a higher probability of cell depolarization.

The effect of cell depolarization (increase on the value of Vmem) on the level of its DNA methylation is introduced by the following equation (like the previous one),

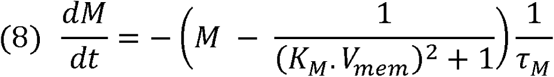

where the methylation level (*M*) increases in time with the decrease of the absolute value of Vmem (depolarized cells have a less negative, or positive, value of Vmem when compared with hyperpolarized ones). The parameter K_M_ gives the inverse of the value of Vmem that decreases the change rate of *M* by 0.5. Again, ⎮_M_ gives the time scale of change in *M*. A decrease of |Vmem|, due to cell depolarization, forces an increase on the stable value of *M* (an increment in ICH methylation). Both previous equations are solved for 100 ms, with time steps of Δt = 0,01 ms.

### Statistics

All statistical analyses were performed using R v4.3.2 (R Core Team, Vienna, Austria) and GraphPad Prism version 5.0.3 for Windows (GraphPad Software, San Diego, California, USA; https://www.graphpad.com). A two-tailed p < 0.05 was considered statistically significant. Normality of continuous variables was assessed using the Shapiro–Wilk test. Parametric tests like Student’s t-Test were used when the data met assumptions of normality and equal variance; otherwise, non-parametric tests such as the Wilcoxon signed-rank test or Welch’s corrected t-Test were applied. For bioelectric measurements expressed as L/R ratios, inter-group differences were assessed using a two-tailed one-sample t-test of the ratio against a theoretical value of 1, since R-sided tumors served as the internal reference for each paired measurement.

For differential DNA methylation analysis, we used edgeR quasi-likelihood F-tests, and genes with FDR < 0.05 were considered significantly differentially methylated. Volcano plots display FDR values against log2 fold-change (L vs R).

For each set of parameters of the computational model, we performed 1000 statistically independent simulation runs. Most of the model parameters values change in each run, following a gaussian distribution about a mean value.

All experiments were performed with at least three biological replicates, each including a minimum of three technical replicates when applicable (e.g., qPCR, Vmem measurements in cultured cells).

## Supporting information

Supplementary enmended

## Data availability

All data generated or analyzed during this study are included in this published article and its supplementary information files and are available at the following URL: https://www.mediafire.com/file/q9xjh6it3t0cu7s/Tabix.tsv.bgz/file

## Code availability

The original datasets generated during the current study (methylation and BAM files) have been deposited in the National Center for Biotechnology Information (NCBI) Gene Expression Omnibus (GEO) repository and are accessible through accession number GSE339572.

Publicly available breast cancer datasets analyzed in this study were retrieved from the following sources:

SCAN-B: The RNA-sequencing gene expression and supplementary datasets were obtained from the Mendeley Data repository at doi.org.

METABRIC: Genomic and clinical annotations were downloaded from cBioPortal (https://cbioportal.org).

TCGA: Genomic and clinical data from The Cancer Genome Atlas breast invasive carcinoma cohort were retrieved from GDC Data Portal.

[15:10, 22/7/2026] Sergio: The following secure token has been created to allow review of record GSE339572 while it remains in private status: qtqnocikjfkhvkd

## Acknowledgements.

We thank Dra. Claudia Lanari (IBYME, CONICET Buenos Aires, Argentina) for providing the PDX tumor model^42^, and Dr. Sergio Benitez (IHEM, CONICET Mendoza, Argentina) for helping in tissue cutting and tumor preparations for the immunofluorescences. We also thank Dr. Lautaro Rivera for helping in Image-J IFI images processing. Finally, we thank our funding agents:

- PICT-2020-03959, National Agency of Scientific and Technical Promotion, Argentina.
- Portugal national funds by FCT - Fundação para a Ciência e Tecnologia, I.P. in the framework of the projects UIDB/04564/2020 and UIDP/04564/2020, with DOI identifiers 10.54499/UIDB/04564/2020 and 10.54499/UIDP/04564/2020, respectively.

## Author contributions

MR conceived the study and designed the experiments. SR performed gene expression experiments, generated mice tumors and analysis by IHC, flow cytometry and confocal microscopy on tissues. SL performed the R-based experiments and analyzed the data. NB performed the flow cytometry experiments on animal tumor cells. OB proposed the idea of using bovine extracts and developed the protocols for PG who performed the conditioned cell culture experiments. DC contributed to the flow cytometry results analyses. JC developed the computational models and performed the data analysis and interpretation. MR, JC, SR and SL wrote the manuscript. Parts of the writing and language editing of this manuscript were assisted by the ChatGPT language model (OpenAI), under the supervision of the authors. All authors read and approved the final version.

