## Supplementary material for "Left-Right Asymmetry in the Breast TME is Associated with Distinct Bioelectric and Epigenetic Tumor Features": Table 1: TABLA 1Fixed.docx

| **Gene Symbol** | **logFC** | **PValue** | **FDR** | **Meth_Status** |
| --- | --- | --- | --- | --- |
| GJA1 | 7,806003077 | 1,57625E-06 | 0,000499673 | Hypermethylated |
| AQP4 | 4,772931414 | 0,002617407 | 0,207429525 | Hypermethylated |
| CACNB3 | 3,459683062 | 0,003276084 | 0,207703737 | Hypermethylated |
| LRRC8C | 2,791388767 | 0,007478103 | 0,263395399 | Hypermethylated |
| CACNG1 | 4,160879993 | 0,037548239 | 0,835873118 | Hypermethylated |
| VDAC2 | 2,844122901 | 0,042189179 | 0,835873118 | Hypermethylated |

| **Gene Symbol** | **logFC** | **PValue** | **FDR** | **Meth_Status** |
| --- | --- | --- | --- | --- |
| HTR3B | -3,700754697 | 0,002067367 | 0,207429525 | Hypomethylated |
| KCNK5 | -3,506235447 | 0,002071773 | 0,207429525 | Hypomethylated |
| TRPV5 | -4,897592398 | 0,004817746 | 0,226547135 | Hypomethylated |
| GABRR3 | -3,853679074 | 0,005353763 | 0,226547135 | Hypomethylated |
| GABRG2 | -4,18362127 | 0,005717278 | 0,226547135 | Hypomethylated |
| HCN1 | -2,690650489 | 0,010278846 | 0,325839408 | Hypomethylated |
| KCNK4 | -2,945352485 | 0,012930558 | 0,372635158 | Hypomethylated |
| SCN3B | -3,301386532 | 0,018640379 | 0,492416682 | Hypomethylated |
| CACNA1F | -2,349115798 | 0,033161452 | 0,808629249 | Hypomethylated |
| KCNA1 | -1,906662241 | 0,040800797 | 0,835873118 | Hypomethylated |
| CLIC1 | -3,539853735 | 0,047043315 | 0,877219466 | Hypomethylated |
