## Supplementary material for "Left-Right Asymmetry in the Breast TME is Associated with Distinct Bioelectric and Epigenetic Tumor Features": Table 2: TABLE 2.docx

| **Parameter Value** | **Runs in each interval of the number of depolarized cells** | | |
| --- | --- | --- | --- |
|  | [0,10] | [11,90] | [91,100] |
| Standard | 891 | 1 | 108 |
| M0=1 | 914 | 3 | 83 |
| M0=0 | 463 | 1 | 536 |
| G0dep=1,9 nS | 953 | 4 | 43 |
| G0dep=2,1 nS | 791 | 2 | 207 |
| G0pol=0,9 nS | 866 | 1 | 133 |
| G0pol=1,1 nS | 888 | 3 | 109 |
| G0max=1,7 nS | 929 | 1 | 70 |
| G0max=1,3 nS | 833 | 11 | 156 |
