## Supplementary enmended for "Left-Right Asymmetry in the Breast TME is Associated with Distinct Bioelectric and Epigenetic Tumor Features"

### Supplementary Figures

#### Supplementary Fig. S1

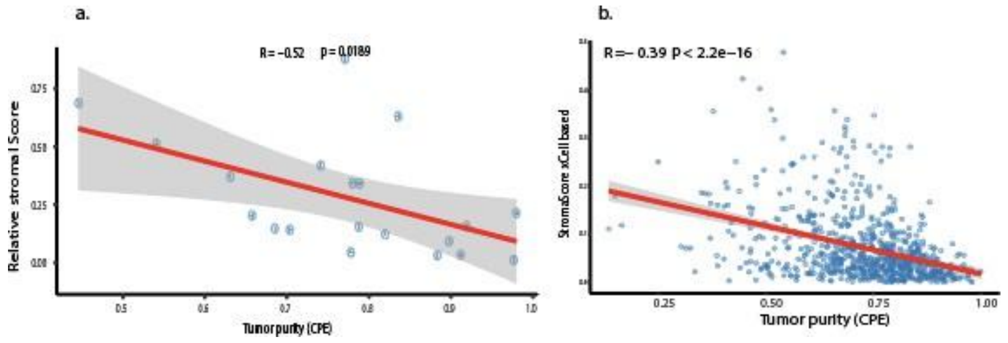


**Supp Figure S1. Correlation between Consensus Purity Estimation (CPE) and stromal abundance in breast IDC samples. a.** Scatterplot showing the relationship between tumor purity (CPE, RNA-Seq–derived) and relative stromal content quantified from 20 H&E-stained TCGA tumor sections using HoVer-Net. A significant negative correlation was observed (Pearson r = –0.519, p = 0.01887), indicating that lower CPE values correspond to higher stromal abundance. **b**. Scatterplot showing the relationship between tumor purity (CPE, RNA-Seq–derived) and stromal content based on RNA-Seq expression data of 141 stromal gene signatures (Pearson r=-0.35, p<0.0001) in 784 TCGA breast IDCs. The red lines represent the regression fit, and the shaded areas the 95% confidence intervals.

#### Supplementary Fig. S2

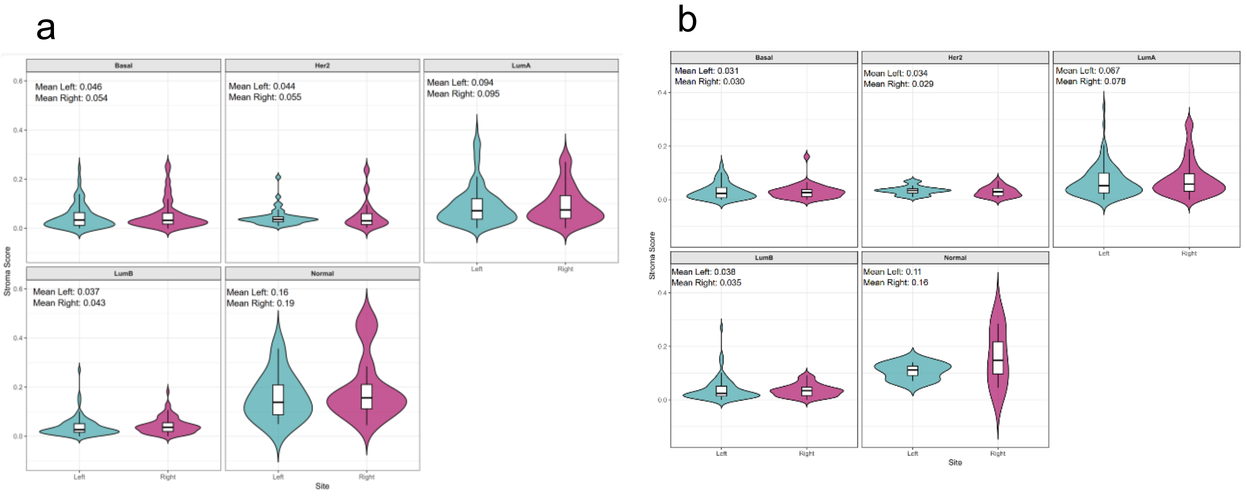

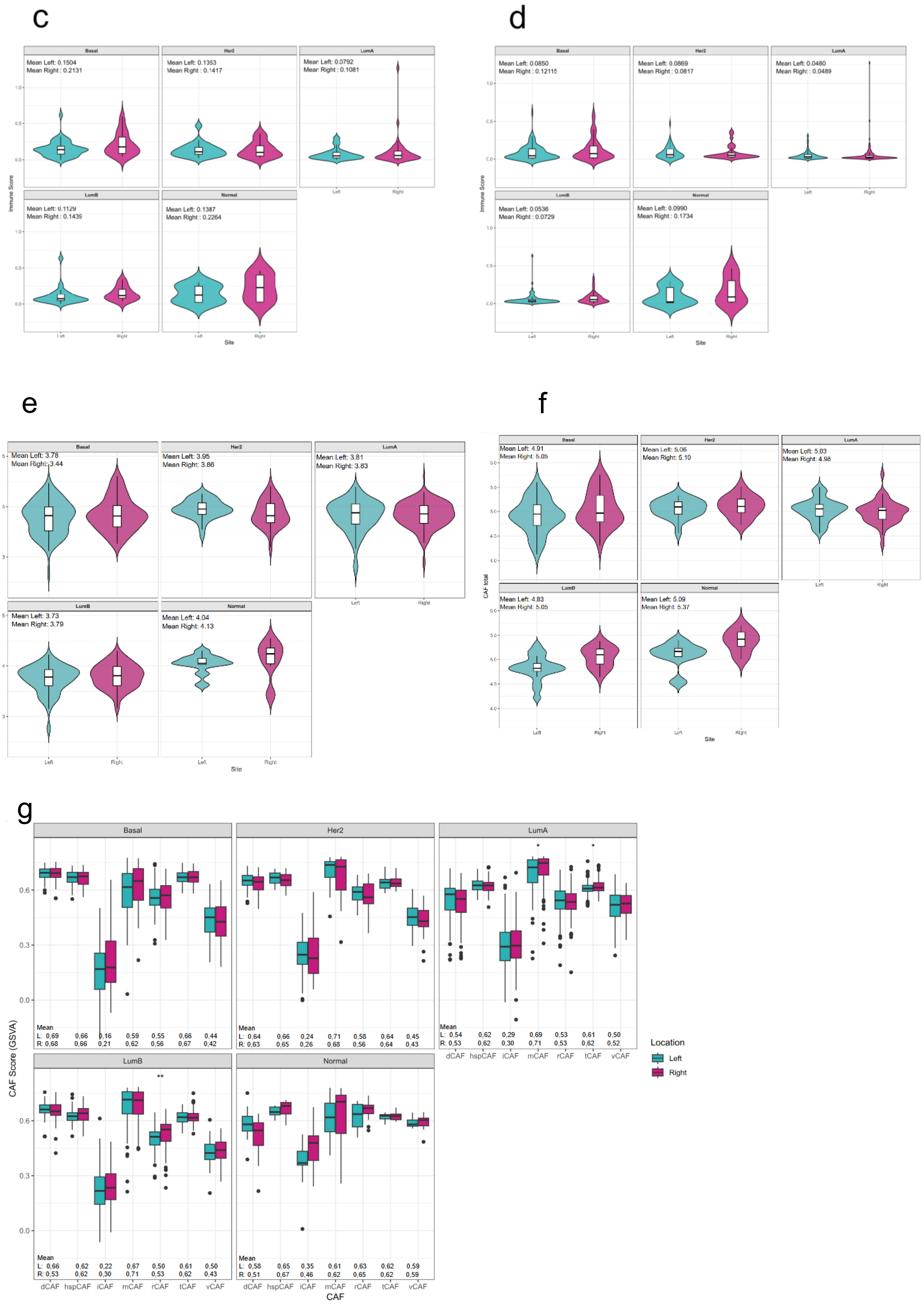


**Supp Figure S2. Stromal composition in L-R breast cancer subtypes calculated with xCell R package.** Given the non-normal distribution of the data, group comparisons were performed using the Wilcoxon rank-sum test. In addition, mean values are reported as descriptive statistics to reflect the overall magnitude of the signature across samples. **a**. Stromal gene signature scores in 5 breast tumor subtypes: basal, HER2, Luminal A, Luminal B and Normal-like from the 276 TCGA IDC cohort with ≤70% purity. The increased R-sided stromal score medians were consistently observed across almost all subtypes although not reaching statistical significance (Wilcoxon test, p>0.05). **b**. The same comparison in the full 784 IDC cohort (no purity restriction). c. Immune gene signature scores across the five breast tumor subtypes (Basal, HER2, Luminal A, Luminal B, and Normal-like) in the 276 TCGA IDC cohort with ≤70% purity. Median scores were directionally higher in right-sided tumors across most subtypes; however, none of these differences were statistically significant (Wilcoxon test, p > 0.05). d. The same comparison in the full 784 IDC cohort (no purity restriction). **e.** Fibroblast gene signature scores in 5 breast tumor subtypes from the 276 TCGA IDC samples (≤70% purity) are elevated in R-tumors of almost all subtypes, although not reaching significance (Wilcoxon test, p>0.05, Fibrooblast_FANTOM_1). **f**. Similar trends in the 784 IDC cohort (no purity restriction). **g.** The L-R differences of dCAFs and iCAFs were consistently observed across almost all PAM50 breast tumor subtypes although not reaching statistical significance (Wilcoxon test, p>0.05)

Supplementary Fig. S3

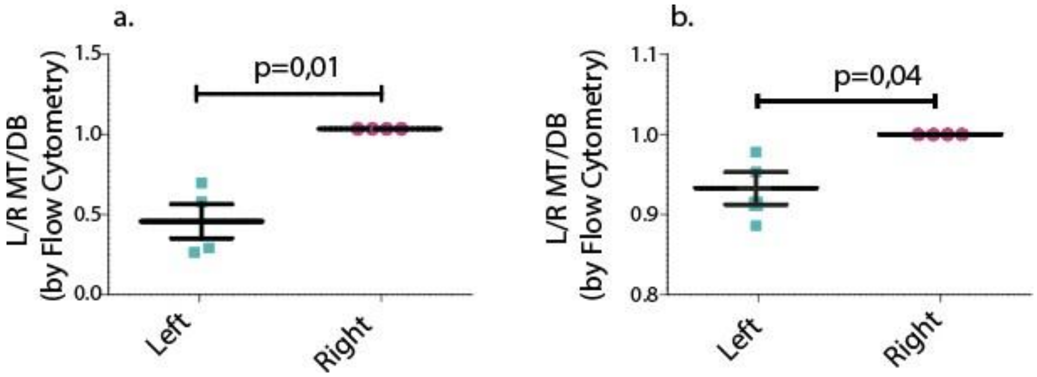


**Supp Figure S3. Membrane potential in L-R conditioned cells.** Cellular Vmem was determined by the ratio between MT and DB fluorescence probes, measured by flow cytometry. Quantifications were first normalized to the mean fluorescence ratio of KCl 65mM treated cells (as a control for maximal depolarization), and results are shown as normalized to the R-treated cells. Cells conditioned with L-side human extracts (panel **a**) and bovine extracts (panel **b**) exhibit a lower MT/DB ratio (depolarized state) compared to those conditioned with R-extracts. Both extract-types separately induce significant differences in the same direction (One-sample t-Test, p<0,05), even though bovine extracts induce smaller differences. Data of L-R are presented as mean ± standard deviation (SD). Analysis performed with GraphPad Prism v5, figure performed with Adobe Illustrator 2024.

Supplementary Fig. S4

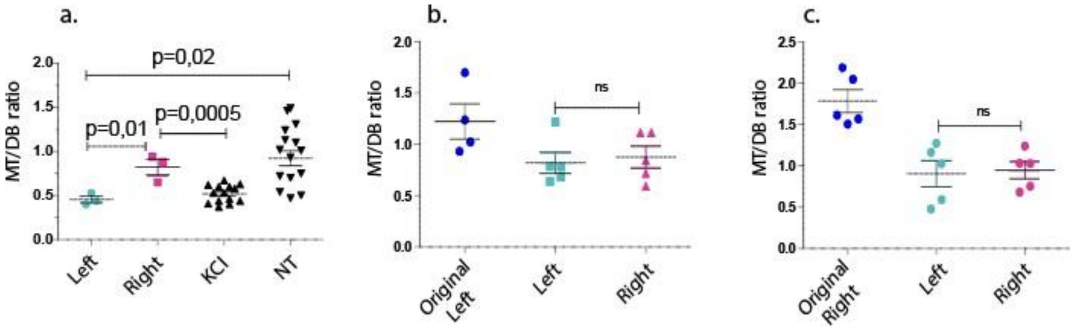


**Supp Figure S4. Vmem in contralateral reimplanted xenografts.** Upon reimplantation of the original tumor fragments into the contralateral mammary glands of new host mice (dark blue: original tumor, panel **a**: original Left, panel **b**: original Right; light blue: reimplanted in left gland, pink: reimplanted in right gland), the L-R Vmem difference was no longer observed, regardless of the tumor's side of origin (paired t-Test, ns). All data are presented as mean ± standard deviation (SD). Analysis performed with GraphPad Prism v5, figure performed with Adobe Illustrator 2024.

Supplementary Fig. S5

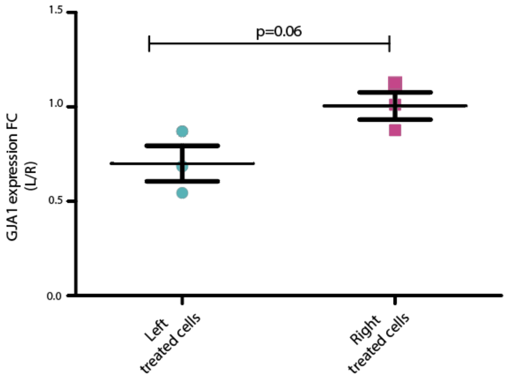


**Supp Figure S5. Gene Expression of Connexins GJA1 in L-R-conditioned MDA-MB231 cells.** Gene expression levels of GJA1 were analyzed in 3 experimental replicates by qPCR. Fold change (FC) is represented as L / R. The quantification shows clear tendency of a reduction in L-treated cells (Unpaired T-test, p=0.06); Data are presented as mean ± standard deviation (SD). Analysis performed with GraphPad Prism v5, figure performed with Adobe Illustrator 2024.

### Supplementary Data

**Supplementary Data 1.** Excel sheet presenting CPE scores and Relative Stromal Abundance of 20 breast IDC samples

**Supplementary Data 2.** Excell sheet presenting gene list for Immune, Stromal and Fibroblast signatures.

**Supplementary Data 3.** Excell sheet presenting gene list of CAF subtypes, based on [31].

**Supplementary Data 4.** Clinical data of 20 H&E-stained whole-slide images of breast IDCs from TCGA.
